# Microglia attenuate regenerative neurogenesis via *sema4ab* after spinal cord injury in zebrafish

**DOI:** 10.1101/2024.10.16.618445

**Authors:** Alberto Docampo-Seara, M. Ilyas Cosacak, Kim Heilemann, Friederike Kessel, Ana-Maria Oprişoreanu, Markus Westphal, Özge Çark, Daniela Zöller, Josi Arnold, Anja Bretschneider, Alisa Hnatiuk, Nikolay Ninov, Catherina G. Becker, Thomas Becker

## Abstract

Zebrafish, in contrast to mammals, regenerate neurons after spinal cord injury, but little is known about the control mechanisms of this process. Here we use scRNA-seq and *in vivo* experiments to show that *sema4ab*, mainly expressed by lesion-reactive microglia, attenuates regenerative neurogenesis by changing the complex lesion environment. After spinal injury, disruption of *sema4ab* doubles the number of newly generated progenitor cells and neurons but attenuates axon regrowth and recovery of swimming function. Disruption of the *plxnb1a/b* receptors, selectively expressed by neural progenitor cells, increases regenerative neurogenesis. In addition, disruption of *sema4ab* alters activation state and cytokine expression of microglia, such that fibroblasts increase expression of the cytokine *tgfb3*, which strongly promotes regenerative neurogenesis. Hence, *sema4ab* in microglia attenuates regenerative neurogenesis in multiple ways, likely directly through *plxnb1a/b* receptors and indirectly, by controlling the inflammatory milieu and *tgfb3* levels. Targeting Sema4A-dependent signalling in non-regenerating vertebrates may be a future strategy to improve regenerative outcomes.

**HIGHLIGHTS:**

- Microglia suppress pro-regenerative fibroblast signalling in a spinal injury site
- Fibroblasts promote regenerative neurogenesis via Tgfb3 signalling
- *sema4ab* promotes microglia activation state after spinal injury
- scRNA-seq reveals full complement of *sema4ab*-dependent changes on different cell types during repair of a spinal lesion site

## INTRODUCTION

Mammals, including humans, fail to regenerate neurons lost to protracted secondary cell death around a spinal injury site (Dusart & Schwab, 1994; Liu *et al*, 1997). Spinal progenitor cells can generate new neurons in a conducive environment and can even be re-programmed *in vivo* to produce oligodendrocytes, but so far not neurons (Llorens-Bobadilla *et al*, 2020; Shihabuddin *et al*, 2000; Stenudd *et al*, 2022). The environment in a spinal injury site consists of immune cells, such as invading blood-derived macrophages (BDMs) and tissue-resident microglia, astrocytes, oligodendrocytes, and fibroblasts. Interactions of these cell types result in an inhibitory environment for regenerative neurogenesis in which prolonged inflammation, as well as fibrotic and astrocytic scarring predominates (Anderson *et al*, 2016; Dias *et al*, 2018; Francos-Quijorna *et al*, 2022; Griffin & Bradke, 2020; Tran *et al*, 2018).

In contrast to mammals, zebrafish functionally regenerate their spinal cord at the adult and larval stage (Becker *et al*, 2004; Chang *et al*, 2021; Mokalled *et al*, 2016; Ohnmacht *et al*, 2016; Vandestadt *et al*, 2021). This includes injury-induced generation of new neurons from resident progenitors (Briona & Dorsky, 2014; Briona *et al*, 2015; Huang *et al*, 2021; Ohnmacht *et al*., 2016; Reimer *et al*, 2008). We use the term ependymo-radial glial cells (ERGs) for these progenitors, by virtue of their combination of ependymal and astroglial features (Becker & Becker, 2015).

A relatively large number of signalling molecules from various cellular sources have been discovered that promote regenerative neurogenesis and axon growth in the zebrafish model (Barreiro-Iglesias *et al*, 2015; Becker & Becker, 2022; Cigliola *et al*, 2023; Goldshmit *et al*, 2012; Mokalled *et al*., 2016; Reimer *et al*, 2009; Reimer *et al*, 2013; Wehner *et al*, 2017). Importantly, regenerative neurogenesis avoids over-proliferation and tumour formation, which may be encountered when transplanting neural progenitor cells to a spinal lesion (Sharp *et al*, 2014), or exhaustion of the progenitor pool, for example observed in a stroke model in mice (Carlen *et al*, 2009). Hence production of new neurons must be precisely regulated in zebrafish, but few negative regulators of regenerative neurogenesis are known in zebrafish (Saraswathy *et al*, 2022).

Microglia are known to promote developmental neurogenesis, e.g. in humans (Yu *et al*, 2025), but activated microglia is mostly detrimental to neurogenesis (Sato, 2015). As microglia signal in the lesion via cytokines, they are in a key position to provide control of neurogenesis. After a spinal cord injury, microglial cells proliferate (Becker & Becker, 2001), but lack expression of the neurogenesis-promoting cytokine *tnfa*, found in BDMs (Cavone *et al*, 2021; Tsarouchas *et al*, 2018), such that their role for regeneration of the spinal cord is unclear. However, in the injured telencephalon, microglia have been found to promote regenerative neurogenesis (Kyritsis *et al*, 2012).

In addition, microglia likely influence the signalling milieu in the entire injury environment (Ewing-Crystal *et al*, 2025), which affects the state of other regeneration-relevant cell types that also influence regenerative neurogenesis. For example, ERGs and astrocyte-like cells produce a range of regeneration-promoting growth factors in central nervous system (CNS) injuries in zebrafish (Cigliola *et al*., 2023; Goldshmit *et al*., 2012; Mokalled *et al*., 2016) and fibroblasts that invade a spinal lesion site promote regeneration by producing favourable extracellular matrix components for axonal regeneration (Kolb *et al*, 2023; Tsata *et al*, 2021; Wehner *et al*., 2017).

To understand how microglia influence the signalling landscape in the injury site and regenerative neurogenesis in zebrafish, we analysed the role of the conserved signalling molecule Semaphorin 4A (Sema4A), which is mainly expressed by microglia and can modulate their activation state (Leitner *et al*, 2015; Meda *et al*, 2012). Sema4A has been shown to promote a pro-fibrotic phenotype in fibroblasts *in vitro*, suggesting possible effects on fibroblasts (Carvalheiro *et al*, 2019). Knock out of the principal Sema4A receptors Plexin B1 and B2 impairs neurogenesis in the developing mouse brain (Daviaud *et al*, 2016), suggesting microglia to progenitor cell communication via Sema4A.

Here, we find that disrupting a zebrafish orthologue of *Sema4A*, *semaphorin4ab (sema4ab)*, increases regenerative neurogenesis, but impairs axon regrowth and functional recovery. *Sema4ab* potentially acts on ERGs via the receptor paralogs *plexin b1a* (*plxnb1a*) and *plexin b1b (plxnb1b*) and indirectly by altering the cytokine milieu, which leads to reduced production of the pro-neurogenic factor *tgfb3* in fibroblasts. Hence, *sema4ab+* microglia regulate regenerative neurogenesis in the regeneration-permissive cellular environment and promote recovery of the zebrafish spinal cord after lesion.

## RESULTS

### *sema4ab*-dependent microglia signalling indicated by scRNA-seq

After a complete spinal lesion at 3 days post-fertilization (dpf), lesion-induced neurogenesis peaks at 24 hours post-lesion (hpl) in larval zebrafish. By 48 hpl, axons have regrown over the lesion site and swimming function is largely recovered (Ohnmacht *et al*., 2016; Wehner *et al*., 2017), such that major events in spinal cord regeneration occur within 48 hours in this system.

To find potential signals from microglia that act on neural progenitors after injury, we interrogated previous scRNA-seq data sets of enriched immune cells and of enriched neural progenitors at 24 hpl in larval zebrafish that had received a spinal cord lesion at 3 dpf (Cavone *et al*., 2021). In the immune cell data set, in which immune cells were enriched by FACS sorting for GFP expression under the *mpeg1.1* regulatory sequences (Suppl. Fig. 1A), we observed a sharp increase in microglia, identified by expression of *apoeb* (Peri & Nusslein-Volhard, 2008), at 24 hpl (Suppl. Fig. 1B, C). We found that the signalling molecule *sema4ab* was expressed by microglia and BDMs (Suppl. Fig. 1D). We searched for potential ligand-receptor interactions with progenitor cells using a data set enriched for expression of *her4.1*, a marker of ERGs, which contain spinal progenitor cells (Suppl. Fig. 1E, F). We found enriched expression of the potential *sema4ab* receptor paralogs *plxnb1a* and *plxnb1b* (Meda *et al*., 2012; Rajabinejad *et al*, 2020) in ERGs (Suppl. Fig. 1G-I). While this indicates the potential for direct ligand-receptor interactions between these two cell types, cell interactions between different types of neural and non-neural cells in the injury site are complex, and altered signalling from microglia is likely to change the entire lesion environment.

To get an overview of most, if not all cells in the injury site and to assess the effect of *sema4ab* ablation simultaneously on all of these cell types, we generated scRNA-seq profiles of the entire trunk tissue flanking the spinal injury site with and without disruption of *sema4ab*. To disrupt *sema4ab,* we injected larvae with highly-active CrRNAs (haCRs) that lead to mutagenesis of the targeted site in nearly all somatic cells, allowing phenotype analysis already in haCR-injected somatic mutants for *sema4ab* (Cavone *et al*., 2021; Hoshijima *et al*, 2019; Keatinge *et al*., 2021; Kroll *et al*, 2021; Saraswathy *et al*, 2024). We generated scRNA-seq profiles of lesion site tissue from 50 control gRNA-injected and 50 *sema4ab* haCR*-*injected larvae at 24 hpl, the peak of regenerative neurogenesis (Ohnmacht *et al*., 2016) (Fig. 1A).

**Fig. 1.**
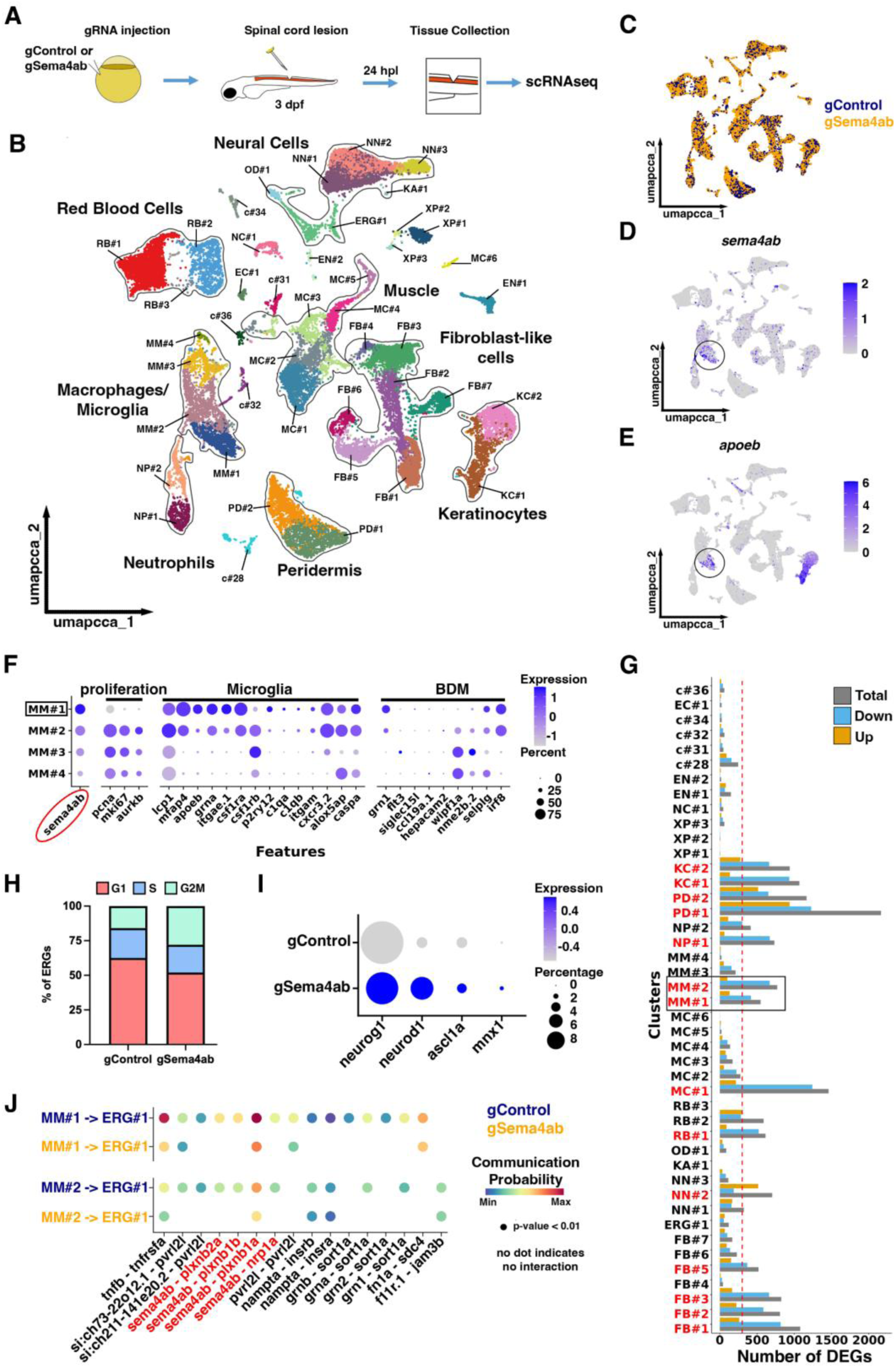
scRNA-seq indicates changes gene expression after *sema4ab* ablation and predicts changes in regenerative neurogenesis. **(A)** A schematic of the experimental design for scRNA-seq in somatic mutants is shown. **(B)** Cluster analysis identifies 44 cell clusters that represent the main expected cell types in the injury site at 24 hpl. **(C)** A UMAP shows no major changes in cell type representation resulting from *sema4ab* disruption (gControl in blue, gSema4ab in yellow). **(D,E)** Feature plots for *sema4ab* (D) and *apoeb* (microglia; E) show overlap (black circles). **(F)** A dot plot of the four BDMs/microglia clusters showing that *sema4ab*-enriched clusters align with the expression of microglia marker genes. **(G)** A graph indicating differential expression of genes (DEG) between samples is shown. Cell clusters with major changes (>300 DEGs) are indicated in red. Microglia clusters MM#1 and MM#2 are boxed. **(H)** Graph shows the proportions of ERGs in each cell cycle phase in control gRNA-injected larvae and *sema4ab* gRNA-injected larvae. Note that the proportion of cells in M/G2 phase is increased in *sema4ab* somatic mutants (gControl: 16% vs gSema4ab: 28%). **(I)** Dot plot showing an increase in differentiation marker gene expression in ERG progenitors when *sema4ab* is disrupted. **(J)** Dot plot illustrating the communication probabilities of all ligand-receptor interactions identified between MM#1 and ERGs, as well as between MM#2 and ERGs. The absence of dots indicates that no interaction was detected. For a key to cluster abbreviations in B and G, see Suppl. table 5.

Our scRNA-seq experiments yielded 11788 cells for the control gRNA-injected sample (gControl), 11621 cells for sample 1 (gSema4abR1), and 11330 cells for sample 2 (gSema4abR2) of two technical repeats of *sema4ab* haCR*-*injected larvae, after quality control (Suppl. Tables 1, 2). The haCR manipulation of *sema4ab* was highly penetrant, as no intact reads of the targeted site could be retrieved, contrary to controls (151 correct reads in controls; 4 and 6 reads, none of which were correct, in *sema4ab* haCR*-*injected larvae; Suppl. Table 3, see Suppl. Table 4 for example reads). Moreover, principal component analysis indicated that the two technical repeats of the *sema4ab* haCR-treated condition were closely related and more distant from the control sample, indicating reproducibility of the approach (Suppl. Fig. 2A).

Unsupervised clustering and annotation revealed 44 distinct cell clusters that fell into 15 major cell categories (Fig. 1B, Suppl. Table 5). Fibroblast-like cells constituted the majority of all cells (21.13 %), consisting of 7 clusters, followed by neural cells (14.86 %), consisting of ERGs, oligodendrocytes and 4 neuronal clusters, red blood cells (13.54 %), consisting of three clusters, muscle cells (13.54 %), consisting of 5 clusters, and immune cells (11.63 %), consisting of 4 BDM/microglia and 2 neutrophil clusters. These clusters were represented in both conditions, as observed in a UMAP projection showing intermingling between cells from control and *sema4ab*-disrupted samples (Fig. 1C). Hence, all major cell types expected from histology were represented in the samples, allowing for comprehensive analyses of cell interactions in the lesioned larval zebrafish spinal cord.

High and selective expression of *sema4ab* was found in a cluster of BDMs and mainly in the *apoeb+* cluster MM#1 (Fig. 1D, E). Using the microglia marker *apoeb* and a marker set from adult zebrafish brain (Zambusi *et al*, 2022), we found that MM#1 and MM#2 expressed higher levels of microglial marker genes, compared to MM#3 and MM#4 that expressed more BDM-related genes. Population MM#2, in contrast to population MM#1, expressed proliferation markers, but otherwise marker expression was very similar between the two populations (Fig. 1F). Therefore, MM#2 likely is a proliferative sub-group of microglia and together with population MM#1 represent microglia. Hence, microglia is likely the main cell type expressing *sema4ab* in the injury site.

To analyse alterations in the scRNA-seq profiles resulting from *sema4ab* disruption, we first compared cell numbers between samples. This indicated no major changes in cell type representation resulting from *sema4ab* disruption with the possible exception of peridermis cluster 2 (PD#2), which showed a ∼50 % reduction in cell number (Suppl. Fig. 2B).

Next, we analysed gene expression changes per cluster. Considerable changes in gene expression (> 300 differentially expressed genes) were present in microglia (MM#1, MM#2), compared to BDM-like clusters (MM#3, MM#4), which correlates with the expression levels of *sema4ab* in these cell cluster. Fibroblasts (FB#1, FB#2, FB#3, FB#5), another regeneration-relevant cell population, also showed high levels of gene expression changes, together with neutrophils (NP#1), neurons (NN#2), muscle (MC#1), peridermal cells (PD#1, PD#2), keratinocytes (KC#1, KC#2) and red blood cells (RB#1) (Fig. 1G). Interestingly, most clusters showed reduced gene expression in the absence of *sema4ab* function, with the exception of the neuronal cluster NN#2 that showed increased gene expression. This indicates changes in gene expression across many cell types when *sema4ab* is disrupted in microglia.

To determine whether such changes may have influenced regenerative neurogenesis, we analysed changes in proliferation and in gene expression related to neuronal differentiation in ERGs. Using the “Cell-Cycling Scoring” analysis code in Seurat (Satija *et al*, 2015) we calculated the proportion of ERGs in different cell cycle phases. We found a substantial increase in ERGs in G2/M phase in sema4ab-injected larvae (gControl: 16%, gSema4ab: 28%) (Fig. 1H). Additionally, a comparison of neuronal differentiation markers (*neurog1*, *neurod1*, *ascl1a*) and motor neuron cell fate marker *mnx1* showed increased expression of these genes in ERGs in the absence of *sema4ab* function (Fig. 1I). This is consistent with enhanced progenitor proliferation and neurogenesis.

To determine potential direct signalling from microglia to ERGs, we analysed altered signal/receptor interactions between microglia and ERGs using CellChat, which evaluates cell interactions based on known ligand-receptor interactions in scRNA-seq data (Jin *et al*, 2021). That indicated reduced interactions of *sema4ab* with *plxnb1a, plxnb1b* and *plxnb2a* (Fig. 1J). Hence, scRNA-seq predicts microglia to directly signal to ERGs using Sema4ab to Plexin receptors.

To get a comprehensive view of how cell types could be affected by changes in microglia, we used CellChat to analyse the outgoing interactions of microglia cluster MM#1. This indicated potentially altered signal/receptor interactions with all identified cell clusters, including ERGs and fibroblasts, but mostly with themselves and their proliferative version MM#2. Most of these signal/ligand interactions were reduced when *sema4ab* was disrupted (Fig. 2A).

**Fig. 2.**
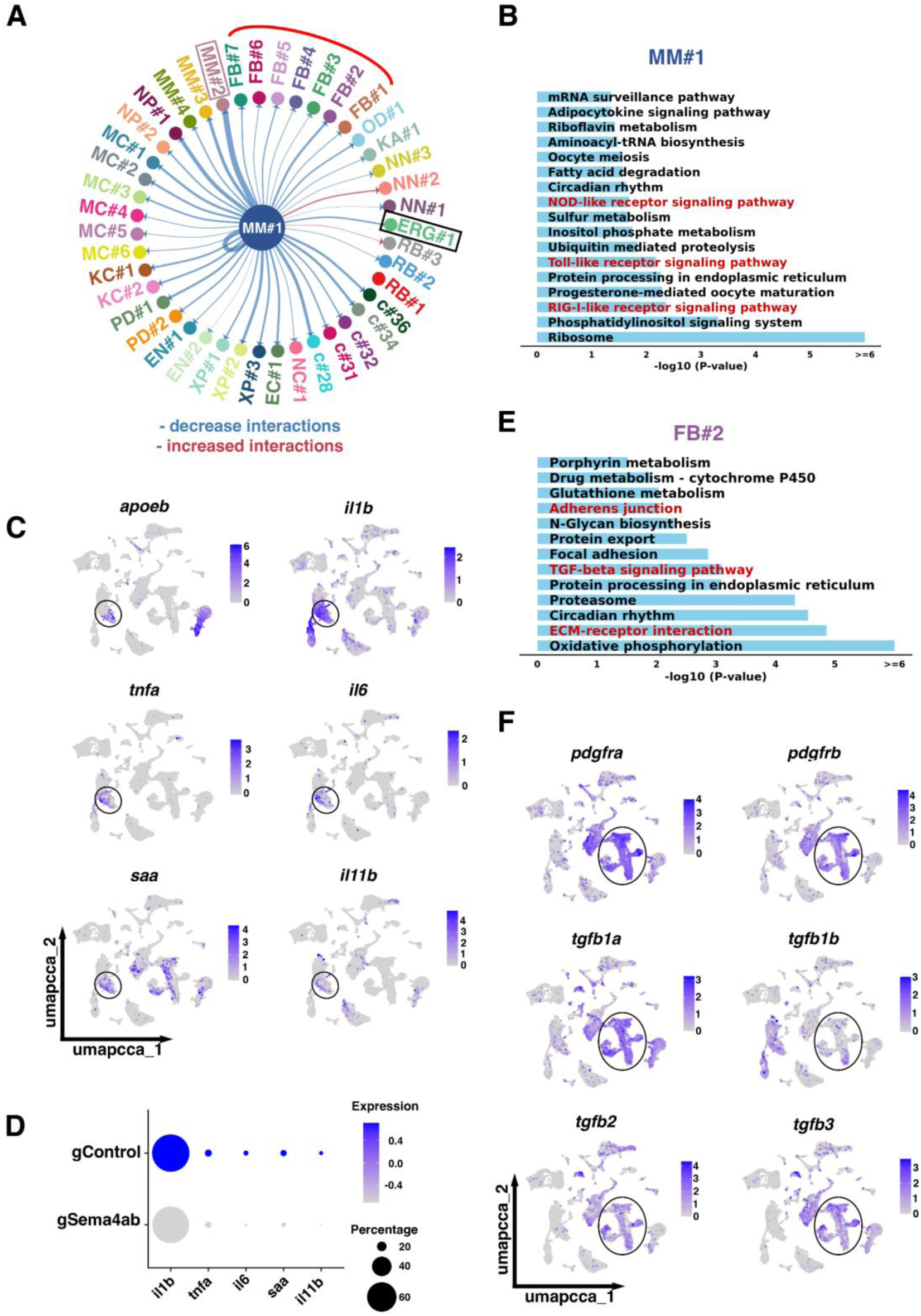
scRNA-seq indicates changes in cell interactions between microglia and other cell types after *sema4ab* ablation. **(A)** CellChat analysis for MM#1 shows interactions with all cell types and a general decrease in these interactions (blue lines) when *sema4ab* is disrupted. Interactions of MM#1 with ERGs (black box), fibroblasts (red curved line) and MM#2 (pink box) are highlighted. **(B)** KEGG analysis of the main cluster that expresses *sema4ab* (MM#1) shows changes in immune activation pathways (highlighted in red). **(C)** Feature plots show enrichment of pro-inflammatory cytokines and inflammatory markers (*il1b, tnfa, il6, saa, il11b)* in microglia (*apoeb*, circle). **(D)** Dot plot showing reduction of expression of pro-inflammatory cytokines and inflammatory markers after *sema4ab* ablation. **(E)** KEGG analysis of the signalling pathways most affected by *sema4ab* disruption in fibroblast-like cell cluster FB#2 is shown. *tgfb* signalling and ECM interaction-related terms are highlighted in red. **(F)** Feature plots showing enrichment of anti-inflammatory cytokines (*tgfb1a, tgfb1b, tgfb2, tgfb3)* in fibroblasts (*pdgfra, pdgfrb;* circled).

To analyse the type of gene expression changes in microglia in the absence of *sema4ab* function, we used GOstats for KEGG analysis (Falcon & Gentleman, 2007) on microglia cluster MM#1. This indicated changes in activation pathways, namely RIG-I-like receptor signalling (Yoneyama *et al*, 2024), Toll-like receptor signalling pathways (Alhamdan *et al*, 2024) and NOD-like receptor signalling (Almeida-da-Silva *et al*, 2023) (Fig. 2B). Most of the indicator genes for these GO terms were under-represented in the sample in which *sema4ab* was disrupted compared to the control sample (20 of 21 genes for RIG-I-pathway, 29 of 32 for Toll-like receptor pathway; 16 of 17 genes for NOD-like receptor signalling; Data file 1), suggesting reduced activation of microglia in the absence of *sema4ab*. Analysis of the top down-regulated genes in MM#1 confirmed this result (Suppl. Table 6). For example, one of the most underrepresented genes after *sema4ab* disruption, *lyz* (5.91-fold down), is related to a pro-inflammatory state (Barros et al., 2015; MacParland et al., 2018). Similarly, *grn1* (6.87-fold down), is linked to a pro-inflammatory state and cytokine production (Horinokita et al., 2019).

Based on this altered activation state of microglia, we analysed expression levels in the scRNA-seq data of common pro-inflammatory cytokines in microglia (MM#1) and indeed found reduced expression for *il1b, tnfa, il6, saa,* and *il11b* (Fig. 2C,D). Reduced activation and pro-inflammatory cytokine production of microglia likely influences different cell types in the injury site.

Of note, fibroblasts have pivotal functions in successful spinal cord regeneration in zebrafish (Tsata & Wehner, 2021), and all 7 clusters of fibroblasts showed reduced incoming signals from microglia, when *sema4ab* was disrupted. Affected signal/receptor interactions included cytokine signalling (*e.g. tnfb, tgfb1b)*, among other pathways (Suppl. Fig. 2C). In addition, KEGG analysis for the three fibroblast populations that showed highest levels of differentially expressed genes after *sema4ab* disruption (FB#1, FB#2, FB#3) indicated ECM-receptor interactions, TGF-beta signalling (enriched in fibroblasts), and adherens junction pathways as changed (shown for FB#2 in Fig. 2E). Interestingly, TGF-beta signalling genes *tgfb1a, tgfb2,* and *tgfb3,* but not *tgfb1b,* are enriched in fibroblasts, compared to BDM/microglia (Fig. 2F) (John et al., 2025). This data suggests that signalling and ECM-related functions were changed in fibroblasts when *sema4ab* was disrupted. Hence, scRNA-seq indicates multiple simultaneous signalling events of *sema4ab+* microglia with other non-neural lesion cells, such as fibroblasts, that likely condition the lesion environment. Overall, scRNA-seq suggests a scenario, in which *sema4ab* signalling from microglial cells to spinal progenitors, as well as via other cell types, attenuates neurogenesis.

### Expression of sema4ab and plexinb1 receptors in vivo

As a first step to verify scRNA-seq results *in vivo*, we determined gene expression patterns of *sema4ab* and *plxnb1a/b* in immune cells and neural progenitors, respectively. We used whole-mounted larvae for multiplexed hybridization chain reaction fluorescence *in situ* hybridization (HCR-FISH) in combination with detection of cell type-specific transgenic markers.

Microglia were identified in *mpeg:mCherry* transgenic fish, in which mCherry is expressed under the *mpeg1.1* promoter (Ellett *et al*, 2011), and by additional expression of *apoeb* (*mpeg:mCherry*+/*apoeb*+). Putative BDMs were *mpeg:mCherry*+/*apoeb*-. In uninjured three-day-old larvae, hardly any *sema4ab*+ cells were detectable, consistent with expression in injury-reactive immune cells. In contrast, at 6 hpl, numbers of *sema4ab*+ cells were strongly increased in the lesion site and remained high to at least 48 hpl (Fig. 3A,B; see Suppl. Fig. 3A for method control). Co-labelling experiments at 24 hpl indicated that the majority of *sema4ab*+ cells were microglia (68.8%; *sema4ab+, mpeg:mCherry+, apoeb+*). 18% were BDMs (*sema4ab*+, *mpeg:mCherry+*, *apoeb-*), and 13.2% were unidentified (*sema4ab*+ only; Fig. 3C-E).

**Fig. 3.**
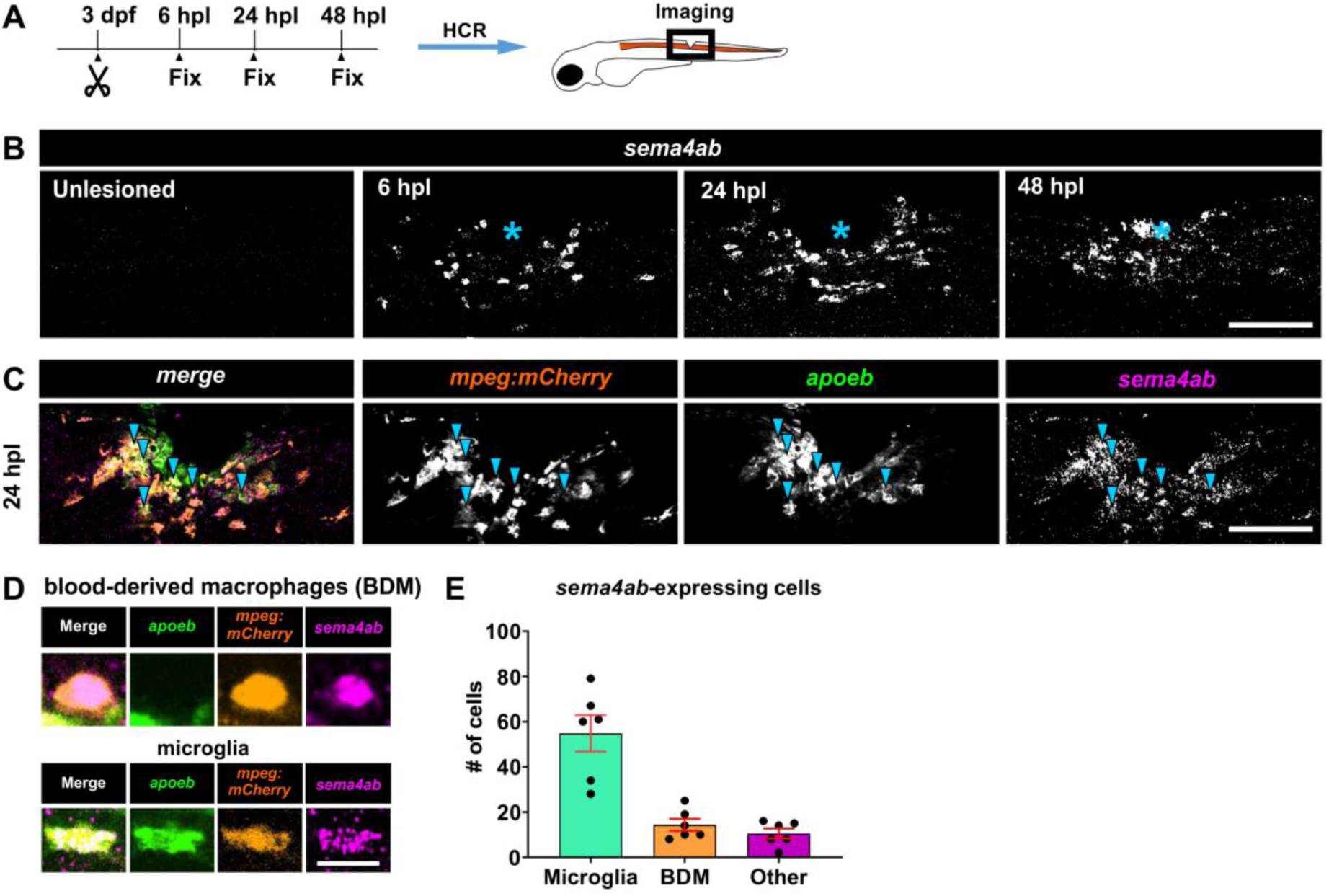
*Sema4ab* is expressed mainly in microglia in the spinal injury site. **(A)** Experimental timeline and schematic of a zebrafish larva indicating the lateral view of the spinal injury site shown throughout the article (unless specified differently) are shown. **(B)** A time-course analysis of HCR-FISH indicates increased expression of *sema4ab* during spinal cord repair. **(C)** Triple-multiplexed HCR-FISH at 24 hpl shows co-localisation of *sema4ab, mpeg:mCherry, and apoeb* at 24 hpl (blue arrowheads). **(D)** High magnifications of *apoeb/mpeg:mCherry/sema4ab* labeling to differentiate blood-derived macrophages (BDMs; *apoeb-*) from microglia (*apoeb+*) for cell counting. **(E)** Quantifications at 24 hpl show that most *sema4ab-*expressing cells are microglia (*apoeb+/mpeg:mCherry+*/*sema4ab+*). Error bars show SEM. Asterisks indicate the injury site. Scale bars: 100 µm (B, C), 15 µm (D).

scRNA-seq analysis from Cavone et al. 2021 showed that *sema4ab* was expressed in neutrophils (see Suppl. Fig. 1D). To determine whether unidentified cells might be neutrophils we determined *sema4ab* expression in neutrophils using the *mpx:GFP* transgenic fish, in which GFP is expressed under the *mpx* promoter (Renshaw *et al*, 2006). Co-labelling of *sema4ab* with *mpx:GFP* expressing neutrophils indicated that 16 % of *sema4ab+* cells were neutrophils at 6 hpl and at 24 hpl (Suppl. Fig. 3B), indicating that the few *sema4ab+* cells not labelled by *mpeg:mCherry* were likely mostly neutrophils. In summary, a spinal lesion induced the presence of *sema4ab+* cells that were mostly microglial cells.

To detect *plexin b1* receptor expresion in vivo, we performed multiplexed HCR-FISH for *plxnb1a* and *plxnb1b* in combination with the *her4.1:EGFP* transgenic reporter line, to label ERGs (Yeo *et al*, 2007). *plxnb1a* mRNA was detectable at low levels in ERGs in unlesioned animals at 3 dpf (Fig. 4A,B). At 24 hpl, the signal appeared slightly weaker, as predicted by lower expression levels in scRNA-seq (see Suppl. Fig. 1G). The signal for *plxnb1b* was generally stronger in ERGs (*her4.1:EGFP+*) than that for *plxnb1a* and did not show detectable changes after lesion (Fig. 4C). High magnifications indicated co-expression of *plxnb1a and plxnb1b* in the same ERGs in unlesioned and lesioned animals. Notably, outside the ERGs only very low (*plxnb1a*) or no signal (*plxnb1b*) was detectable in spinal tissue (Fig. 4D). In the non-neural lesion site, BDMs/microglia (*mpeg:mCherry*+) were also negative for *plxnb1a and plxnb1b* (Fig. 4E). This indicated that transcripts for the paralogous *plxnb1a* and *plxnb1b* were enriched in ERGs, as predicted by scRNA-seq (see Suppl. Fig. 1I).

**Fig. 4.**
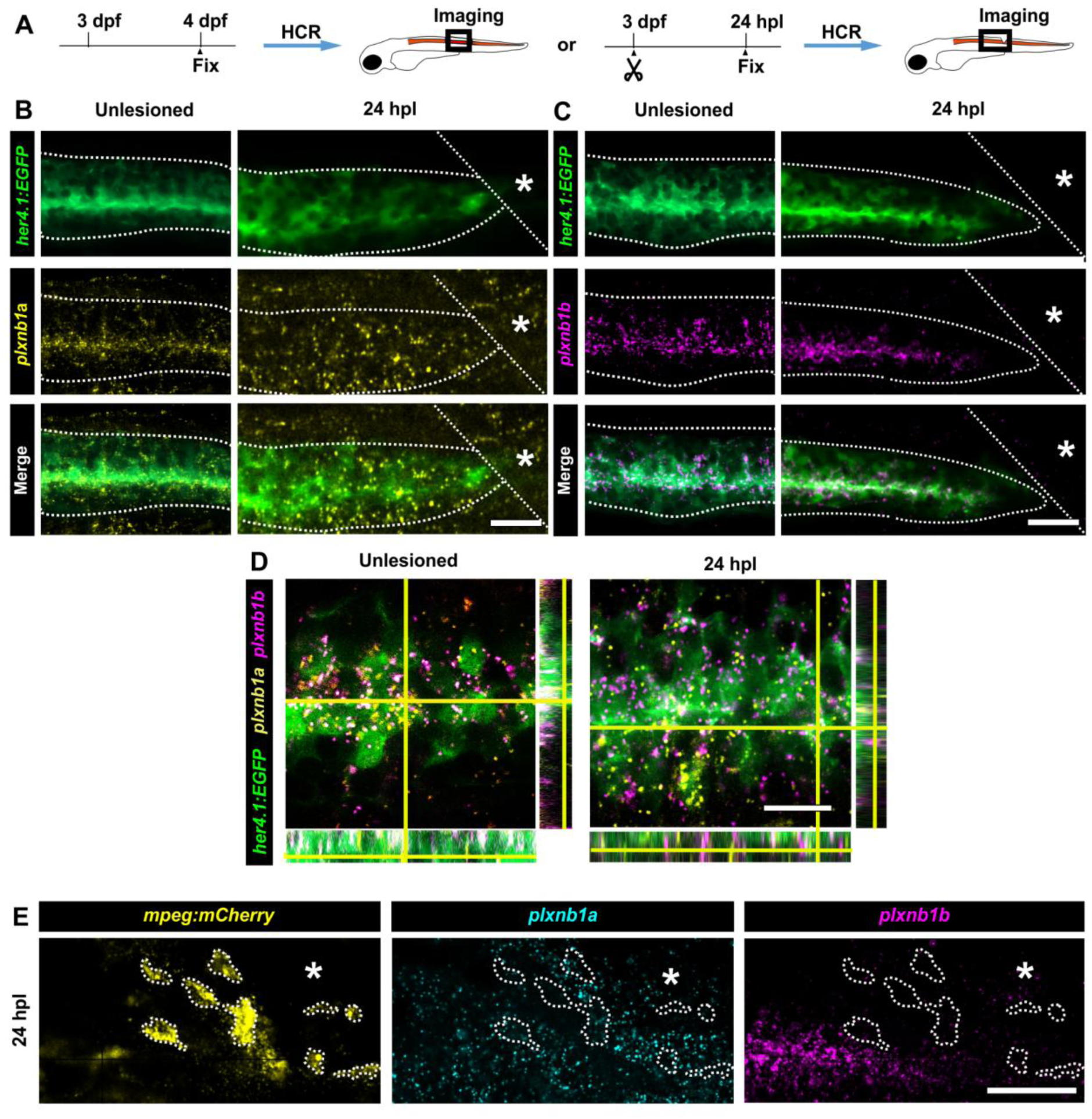
*plxnb1a* and *plxnb1b* are selectively expressed by spinal progenitors. **(A)** A schematic indicating the experimental design for HCR. Photomicrographs are oriented as in Fig. 3. **(B-C)** HCR-FISH in the *her4.1:EGFP* line indicate expression of *plxnb1a* (B) and *plxnb1b* (C) in spinal progenitor cells (*her4.1:EGFP*+) before and after injury. Dotted lines indicate the spinal cord boundaries. **(D)** High magnifications with orthogonal views of co-localization of *plxnb1a* and *plxnb1b* in *her4.1:EGFP*+ cells before and after injury are shown. **(E)** HCR-FISH in the *mpeg:mCherry* line indicates very low expression of *plxnb1a* and no detectable expression of *plxnb1b* in BDM/microglial cells (outlined) after injury. Asterisks indicate the injury site in (B,C,E). Scale bars: 50 µm (B, C, E) 10 µm (D).

### *sema4ab* disruption increases regenerative neurogenesis

For a loss-of-function analysis of *sema4ab* in regenerative neurogenesis, we targeted *sema4ab* in somatic mutants by injecting haCRs into fertilised eggs. We controlled each experiment by an internal control RFLP to show >90% effectiveness of guides in destroying a targeted restriction enzyme recognition site (Suppl. Fig. 4A). Further evidence for the effectiveness of gene disruption comes from the complete disappearance of correct DNA sequences in scRNA-seq analysis (see above and see Suppl. Tables 3, 4). Potential stress responses to the injection are controlled for by always using control gRNA injections in internal controls for each experiment (Johnston *et al*, 2020). Somatic targeting reduces mRNA detectability of *sema4ab* in qRT-PCR by 50%, indicating that gene disruption perturbed mRNA transcription (Suppl. Fig. 4A). Finally, we raised a germline stable mutant for *sema4ab* to verify gene disruption phenotypes. The germline mutant introduced a 14 bp insertion in exon 6, leading to a premature stop codon, which is predicted to lead to a truncated protein lacking functionally important domains, such as sema superfamily domains, plexin repeat domains and the intracellular part, containing a sema4F domain (Suppl. Fig. 4B).

Somatic mutants for *sema4ab* showed no differences in growth compared to control gRNA injected larvae, as measured by body length (Suppl. Fig. 4C-E), notochord thickness (Suppl. Fig. 4F), and spinal cord thickness (Suppl. Fig. 4G). However, a small decrease in eye size was observed (8 % reduction, Suppl. Fig. 4H). Germline mutants showed no change in body length (Suppl. Fig. 4I-K) and replicated the reduction in eye diameter seen in somatic mutants (9 % reduction; Suppl. Fig. 4L). Slightly reduced eye size may be related to compromised photoreceptor maintenance in the absence of Sema4a function (Toyofuku *et al*, 2012). Hence, *sema4ab* disruption was effective in somatic and germline mutants and gross development of larvae was largely unimpaired in the absence of *sema4ab*.

To analyse regenerative neurogenesis in somatic and germline stable mutants, we used 5-Ethynyl-2’-deoxyuridine (EdU) labelling in the *mnx1:GFP* transgenic line that indicates motor neurons (Flanagan-Steet *et al*, 2005). Motor neurons represent the main cell type that is newly generated during lesion-induced neurogenesis (Cavone et al., 2021). In unlesioned *sema4ab* haCR-injected somatic mutant larvae (EdU added at 72 hpf, analysis at 96 hpf) few newly generated motor neurons (EdU+/*mnx1*:*GFP*+) were present, similar to control gRNA-injected larvae (Fig. 5A). This was expected, since *sema4ab* was not detectable in the spinal cord without lesion and shows that *sema4ab* does not have a detectable role in ongoing low-level developmental neurogenesis.

**Fig. 5.**
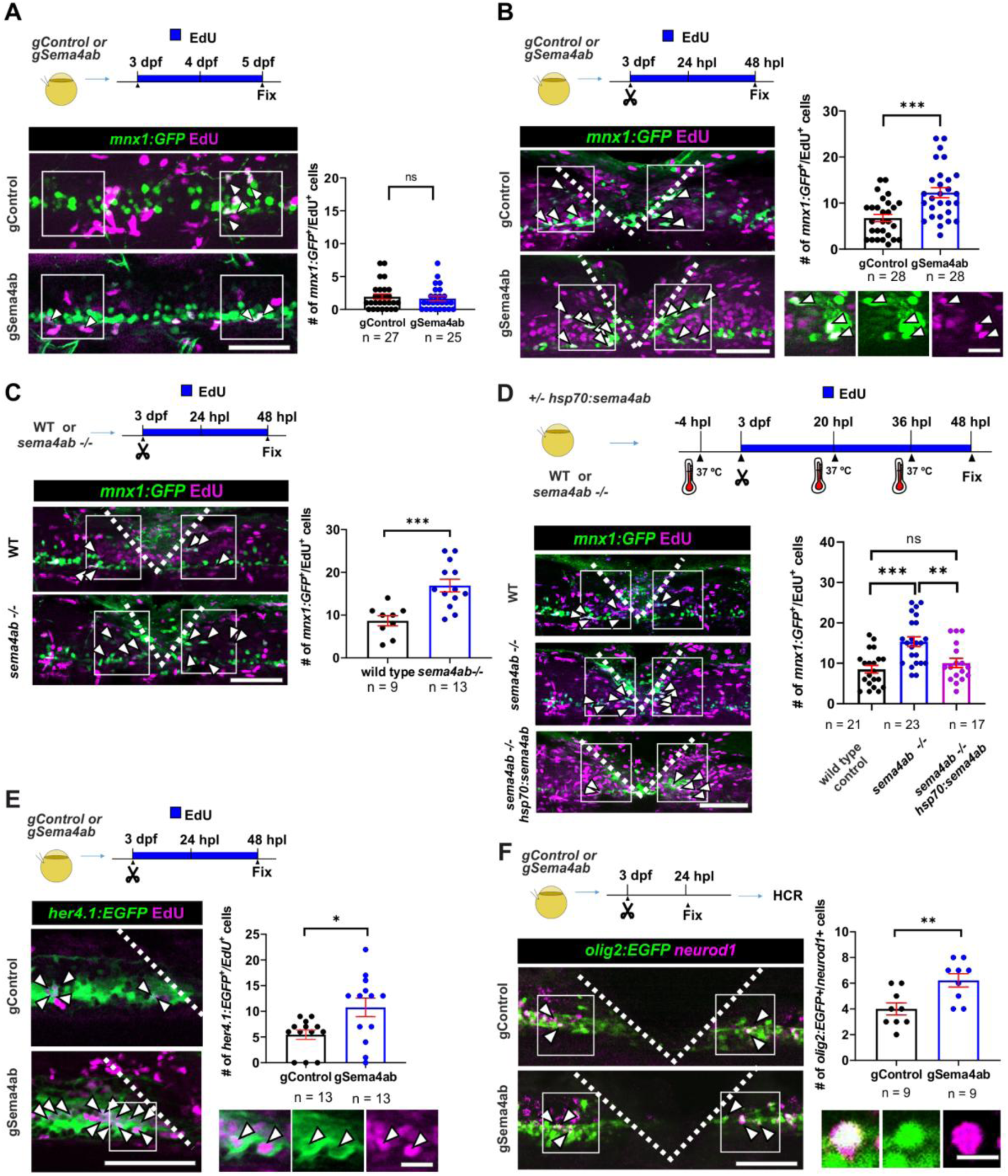
*Sema4ab* loss of function increases regenerative neurogenesis. **(A)** Ongoing developmental neurogenesis of motor neurons (*mnx1:GFP*+/EdU+) is not different between gControl- and gSema4a-injected unlesioned larvae (gControl: 1.9 cells per larvae ± 0.3920; gSema4ab: 1.7 cells per larvae ± 0.3646; Mann-Whitney U-test: p = 0.5972). **(B)** Lesion-induced generation of motor neurons (*mnx1:GFP*+/EdU+) is increased in somatic mutants for *sema4ab* (gControl: 6.9 cells per larva ± 0.8; gSema4ab: 12.9 cells per larva ± 1.1; Mann-Whitney U-test: p = 0.0001). (**D)** Lesion-induced generation of motor neurons (*mnx1:GFP*+/EdU+) in germline mutants for *sema4ab* is also increased (wild type: 8.7 cells per larva ± 1.2; *sema4ab*-/-: 16.9 cells per larva ± 1.5; Mann-Whitney U-test: p = 0.0001). **(D)** Overexpression of *sema4ab* driven by a heat-shock promotor (*hsp70:sema4ab*) reduces over-production of motor neurons (*mnx1:GFP*+/EdU+) back to wild type levels in *sema4ab* germline mutants (wild type: 8.5 cells per larva ± 0.95; *sema4ab*-/- : 15.4 cells per larva ± 1.2; *sema4ab*-/- + *hsp70:sema4ab*: 10 cells per larva ± 1.15; Kruskal-Wallis test: p = 0.0004; Dunn’s multiple comparisons test: ***p = 0.006; n.s > 0.9999, **p = 0.0168). All conditions shown were heat-shocked. **(E)** Lesion-induced proliferation of ERGs (*her4.1:EGFP+*/*EdU+*) is increased in somatic mutants for *sema4ab* (gControl: 5.4 cells per larva ± 0.9; gSema4ab: 10.77 cells per larva ± 1.8; Mann-Whitney U-test: p = 0.0219). **(F)** The number of *neurod1*-labelled motor neuron progenitor cells (*olig2:EGFP+/neurod1+*) is increased after *sema4ab* disruption (gControl: 4.0 cells per larvae ± 0.4714; gSema4ab: 6.2 ± 0.5212 cells per larvae; Unpaired t-test: p= 0.0060). Error bars show SEM. Scale bars: 50 µm (A, B, C, D, E, F), 10 µm (insets).

After a lesion at 3 dpf, we observed a ∼4-fold lesion-induced increase in the number of new motor neurons in control gRNA-injected larvae in the vicinity of the lesion at 48 hpl, as previously reported (Ohnmacht et al., 2016). Remarkably, in *sema4ab* haCR injected larvae, this number was further increased by 85 %, compared to lesioned wild type control gRNA-injected larvae (Fig. 5B). Similarly, in the lesioned *sema4ab* germline mutant, the number of new motor neurons almost doubled (95 % increase) compared to lesioned wild type control larvae (Fig. 5C). This result is in agreement with the prediction by scRNA-seq of increased neurogenesis after *sema4ab* disruption.

To determine whether this phenotype is likely due to *sema4ab* loss-of-function, we over-expressed the protein in *sema4ab* germline mutants to see a possible rescue effect. To that aim, we generated a construct that leads to marked overexpression of *sema4ab* and a fluorescent reporter under the control of a heat-shock promotor (*hsp70l:sema4ab-t2a-mCherry,* abbreviated as *hsp70:sema4ab)* in heat-shocked plasmid-injected, but not in control animals (Suppl. Fig. 5A-D). Indeed, over-expression in the *sema4ab* germline mutant reduced the number of newly generated motor neurons to that observed in wildtype controls (Fig. 5D), indicating a phenotypic rescue. Over-expression of *sema4ab* in wildtype animals did not alter the number of newly generated motor neurons (Suppl. Fig. 5E), which may indicate a ceiling effect for *sema4ab* expression. The phenotypic rescue in the *sema4ab* mutant confirmed that the germline mutant carried a *loss-of-function* mutation, and that *sema4ab* is a strong negative regulator of regenerative neurogenesis.

To determine how the increased number of new motor neurons observed after *sema4ab* disruption was related to progenitor proliferation, we cumulatively labelled *her4.1:GFP*-expressing ERGs with EdU (Fig. 5E). This indicated a doubling (97% increase) in EdU-labelled ERGs (*her4.1:GFP*+/EdU+) in lesioned *sema4ab* somatic mutant larvae, compared to control-gRNA injected lesioned larvae. This suggests that *sema4ab* negatively affects progenitor proliferation.

To determine whether there was a change in the number of ERGs undergoing neuronal differentiation, we analysed the expression of the proneural transcription factor *neurod1* (Pataskar et al, 2016) in motor neuron progenitor cells by HCR-FISH. The number of *neurod1*-expressing motor neuron progenitor cells (*olig2:EGFP+/neurod1+*) was increased by 50% in lesioned somatic mutants for *sema4ab*, compared to lesioned control-gRNA injected larvae, at 24 hpl (Fig. 5F). Hence, in the absence of *sema4ab* function, the number of EdU label-retaining progenitor cells, the number of neurogenic progenitors, and newly generated neurons were all increased. This is consistent with scRNA-seq data and indicates that in unperturbed regeneration, *sema4ab* restricts proliferation of neurogenic progenitors to reduce regenerative neurogenesis.

### *sema4ab* disruption impairs recovery of swimming function and axon regrowth

Behavioural recovery is an important aspect of spinal repair. Therefore, we assessed functional recovery by measuring the distance swum after a vibration stimulus. Uninjured *sema4ab* germline mutants swam distances that were not different from those of uninjured wild type larvae at 5 dpf (Fig. 6A). In contrast, after a previous spinal cord transection at 3dpf, we observed a 35 % reduction in swim distance in lesioned *sema4ab* germline mutants, compared to lesioned wild type animals (Fig. 6B).

**Fig. 6.**
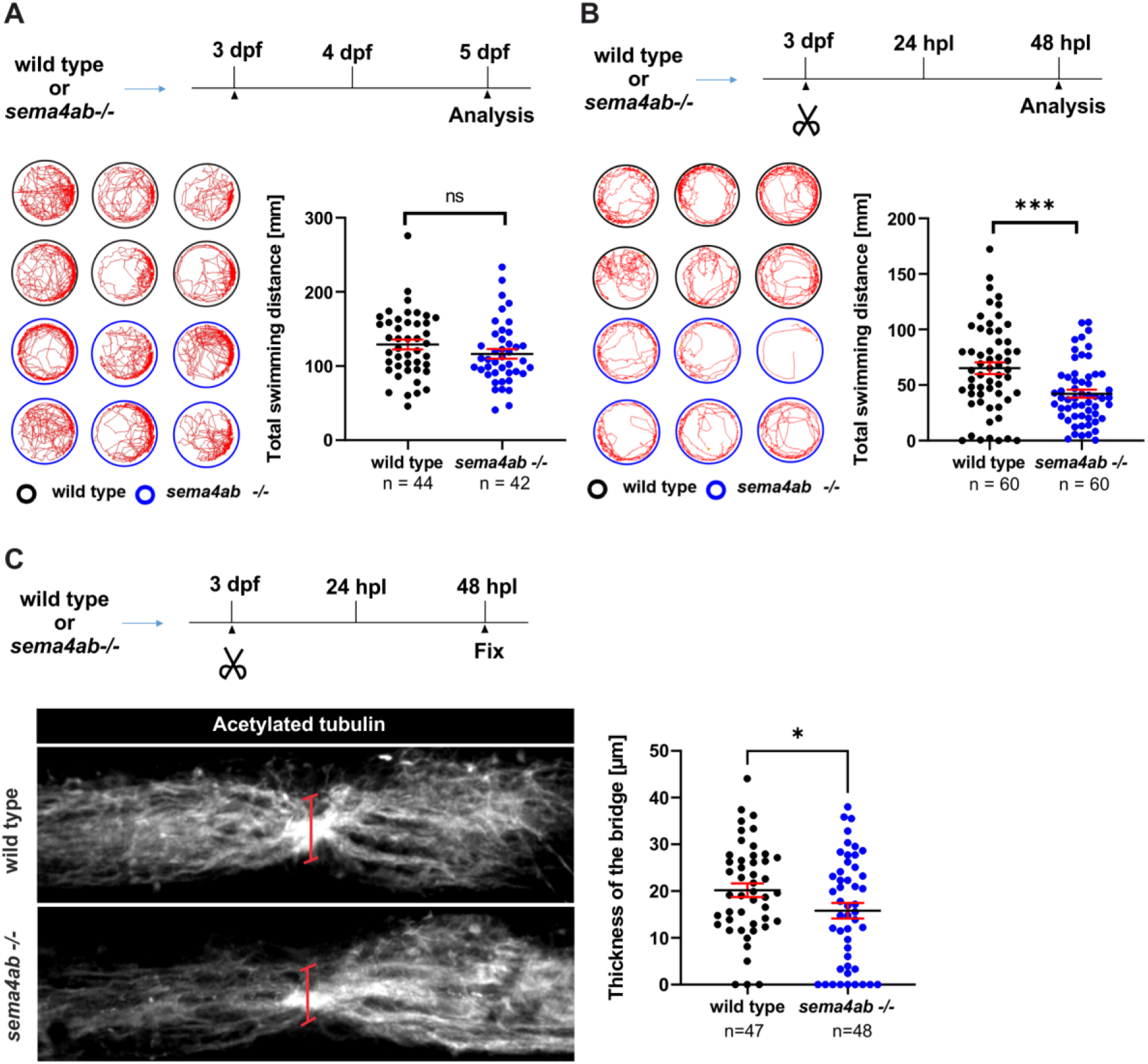
Disruption of *sema4ab* impairs recovery of swimming behaviour. **(A)** A test of swimming behaviour at 5 dpf in unlesioned larvae shows no change in swimming distance in *sema4ab* germline mutants, compared to wildtype controls (wild type: 128.8 mm/min ± 6.657; *sema4ab* -/-: 116.2 mm/min ± 6.504; Unpaired t-test: p = 0.1798). **(B)** A test of swimming behaviour at 48 hpl in 5 dpf larvae shows a reduction in swimming distance in *sema4ab* germline mutants (wild type: 76.98 mm/min ± 8.846; *sema4ab* -/-: 42.15 mm/min ± 5.508; Mann-Whitney U-test: p = 0.0005). **(C)** In *sema4ab* mutants the axonal fibre bundle crossing the injury site shows reduced thickness, compared to controls at 48 hpl (wild type: 20.20 µm ± 1.5; *sema4ab* -/-: 15.8 µm ± 1.7; one-tailed t-test: p = 0.0253). Error bars show SEM. Scale bars: 50 µm.

Since recovery from spinal injury strongly correlates with axon regrowth across the lesion (Wehner *et al*., 2017), we tested whether axon regrowth would be reduced in *sema4ab* mutants. We assessed the thickness of the neuronal tissue bridge across the lesion by immunolabelling of axons re-connecting the spinal cord stumps in germline mutants for *sema4ab* (Fig. 6C) (Wehner *et al*., 2017). Indeed, we observed a 25 % reduction in the thickness of the axonal bridge at 48 hpl in lesioned *sema4ab* germline mutant animals, compared to lesioned wild type control animals. These results demonstrate that *sema4ab* does not detectably affect development of the locomotor system, but is necessary for efficient recovery of swimming function and axon regrowth after injury.

### Disruption of *plxnb1a/b* augments regenerative neurogenesis

Enriched expression of *plxnb1a* and *plxnb1b* in ERGs after injury made these good candidates for receiving a *sema4ab* signal. Therefore, we disrupted *plxnb1a* and *plxnb1b* by haCR injection. Internal RFLP controls confirmed high efficiency of the haCRs and mRNA expression was reduced by 55-60% for these genes (Suppl. Fig. 6A). While single somatic mutations had no effect (Fig. 7A, B), simultaneous mutation of *plxnb1a* and *plxnb1b* by co-injecting the respective haCRs increased numbers of newly generated motor neurons (EdU+/*mnx1:GFP*+) by 55 % at 48 hpl (Fig. 7C). Disruption of *plxnb2b*, which was also expressed in ERGs, and its paralog *plxnb2a* (Suppl. Fig. 6B-E), had no effect in regenerative neurogenesis (Suppl. Fig. 6F-H). Hence, disrupting *plxnb1a/b*, which code for Sema4ab receptors and are enriched in ERGs, only partially replicated the effect of *sema4ab* mutation. This indicates that while some direct signalling from microglia to ERGs is likely, there is probably also a major effect from the changed lesion site environment after *sema4ab* disruption.

**Fig. 7.**
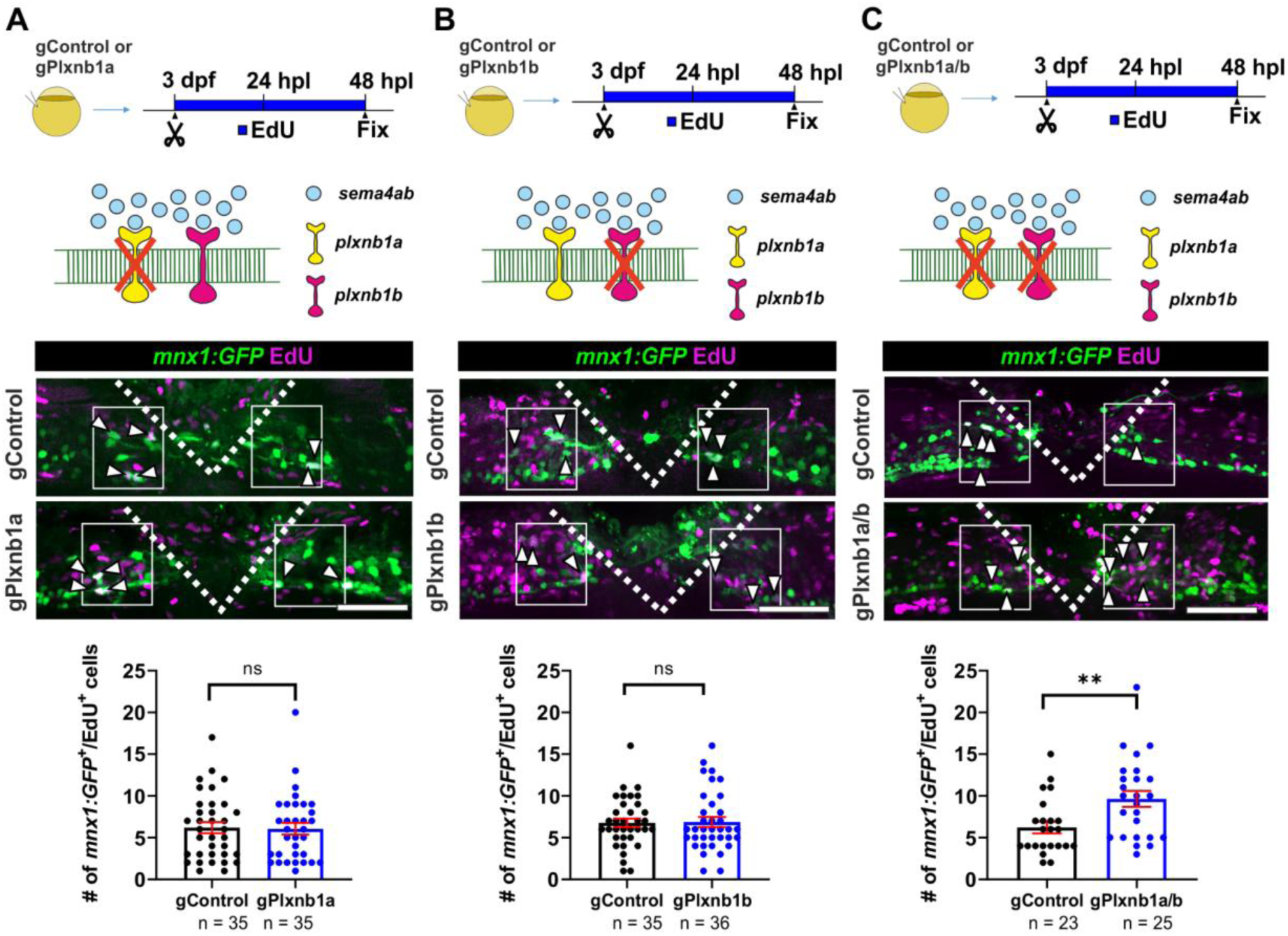
Combined *plxnb1a* and *plxnb1b* loss of function increases regenerative neurogenesis. Photomicrographs are oriented as in Fig. 2. **(A,B)** Regenerative neurogenesis is not changed in single somatic mutants for *plxnb1a* (A, gControl: 6.2 cells per larva ± 0.66 ; gPlxnb1a: 6.1 cells per larva ± 0.71; Mann-Whitney U-test: p = 0.8371) and *plxnb1b* (B, gControl: 6.8 cells per larva ± 0.52 ; gPlxnb1b: 6.9 cells per larva ± 0.60; Mann-Whitney U-test: p = 0.6511). **(C)** Regenerative neurogenesis is increased in double somatic mutants of *plxnb1a and plxnb1b* (gControl: 6.2 cells per larva ± 0.71 cells per larva; gPlxnb1a/b: 9.6 cells per larva ± 0.96; Mann-Whitney U-test: p = 0.0057). Error bars show SEM. Scale bars: 50 µm

### Disruption of *sema4ab* reduces the number of microglia but does not change numbers of other invading cell types

To characterize the influence of *sema4ab* on the lesion site environment *in situ*, we analysed the numbers of cells invading the lesion site with known roles in regeneration, namely neutrophils, fibroblast-like cells, microglia, and BDMs (de Sena-Tomas *et al*, 2024; Tsarouchas *et al*., 2018; Tsata *et al*., 2021; Wehner *et al*., 2017). The number of neutrophils (*mpx:GFP*+) in the lesion site was unchanged between lesioned control gRNA-injected larvae and *sema4ab* haCR-injected larvae at 4 hpl (peak of neutrophil invasion) and 24 hpl (peak of regenerative neurogenesis; Suppl. Fig. 7A-C). The density of fibroblasts (*pdgfrb:GFP+*) also remained unchanged in the absence of *sema4ab* function at 24 hpl (Suppl. Fig. 7D).

For BDMs and microglia (*mpeg*:*mCherry*+) we observed a moderate reduction in number by 12 % in *sema4ab* haCR injected larvae (Suppl. Fig. 7E) at 24 hpl. To determine whether microglia and BDMs would be differently affected in the absence of *sema4ab*, we separately quantified BDMs and microglia by co-labelling with the 4C4 microglia marker antibody (Becker & Becker, 2001). This showed that BDMs (*mpeg:mCherry+;* 4C4-) did not change in cell numbers, whereas microglia numbers (*mpeg:mCherry+;* 4C4+) were reduced by 25% (Suppl. Fig. 7F).

We also assessed potential changes in proliferation of BDMs and microglia using cumulative EdU labelling in combination with immunohistochemistry with 4C4 labelling in the *mpeg:mCherry* reporter line (Suppl. Fig. 7F). This showed that 44% of BDMs and 57% of microglia were newly generated in control animals after injury. The number of newly generated BDMs (*mpeg:mCherry+*, EdU+, 4C4-) was not changed in *sema4ab* somatic mutants after lesion. In contrast, the number of newly generated microglia (*mpeg:mCherry+;* EdU+, 4C4+) was reduced by 33%. Hence, microglia numbers were reduced, likely related to impaired proliferation in the absence of *sema4ab*.

We next analysed phagocytosis of cellular debris, which is one of the major behaviours of BDMs/microglia (Yu *et al*, 2022). We used acridine orange staining to detect levels of cellular debris as a proxy for efferocytosis. The average fluorescence intensity of acridine orange in the lesion site was comparable between control gRNA-injected larvae and *sema4ab* somatic mutants at 24 hpl (Suppl. Fig. 7G). Similarly, the number of BDMs/microglia with acridine orange-positive inclusions, as indicators of phagocytosed debris, did not show differences in these experimental groups (Suppl. Fig. 7H). This indicates that efferocytosis of debris by BDMs/microglia was not quantitatively altered in *sema4ab-*deficient animals.

### Cytokine signalling is altered in *sema4ab*-deficient animals

Changes in cytokine levels, as indicated by our scRNA-seq analysis, can have profound effects on cell behaviour. Therefore, we analysed expression levels of *serum amyloid A* (*saa*), an inflammation marker and regulator of cytokine release (Song *et al*, 2009), and major pro-(*il1b, tnfa, il6, il11b*) and anti-(*tgfb1a, tgfb3*) inflammatory cytokines (Woodcock & Morganti-Kossmann, 2013) in the lesion site environment in control gRNA- and *sema4ab* haCR-injected lesioned larvae by qRT-PCR at 4 and 24 hpl (Fig. 8A). *il1b*, *tnfa*, and *il6* did not show any changes at 4 hpl (Fig. 8B), when the immune response was dominated by neutrophils. However, reduced expression levels of pro-inflammatory cytokines *il1b* (by 50 %), *tnfa* (by 75 %), and *il6* (by 65 %), together with those of *il11b* (by 45 %) and *saa* (by 75 %), were detected at 24 hpl (Fig. 8C), when microglia dominated the injury site. At that time point, levels of *tgfb1a* expression were slightly reduced (by 20 %), whereas those of *tgfb3* were more than doubled (120 % increase; Fig. 8D). In histology, fluorescence intensity for *il1b:GFP* and *tnfa:EGFP* was likewise reduced in *sema4ab* somatic mutants (Suppl. Fig. 8A-C). Overall, these results indicate a shift from a pro-inflammatory to an anti-inflammatory lesion environment in the absence of *sema4ab* function.

**Fig. 8.**
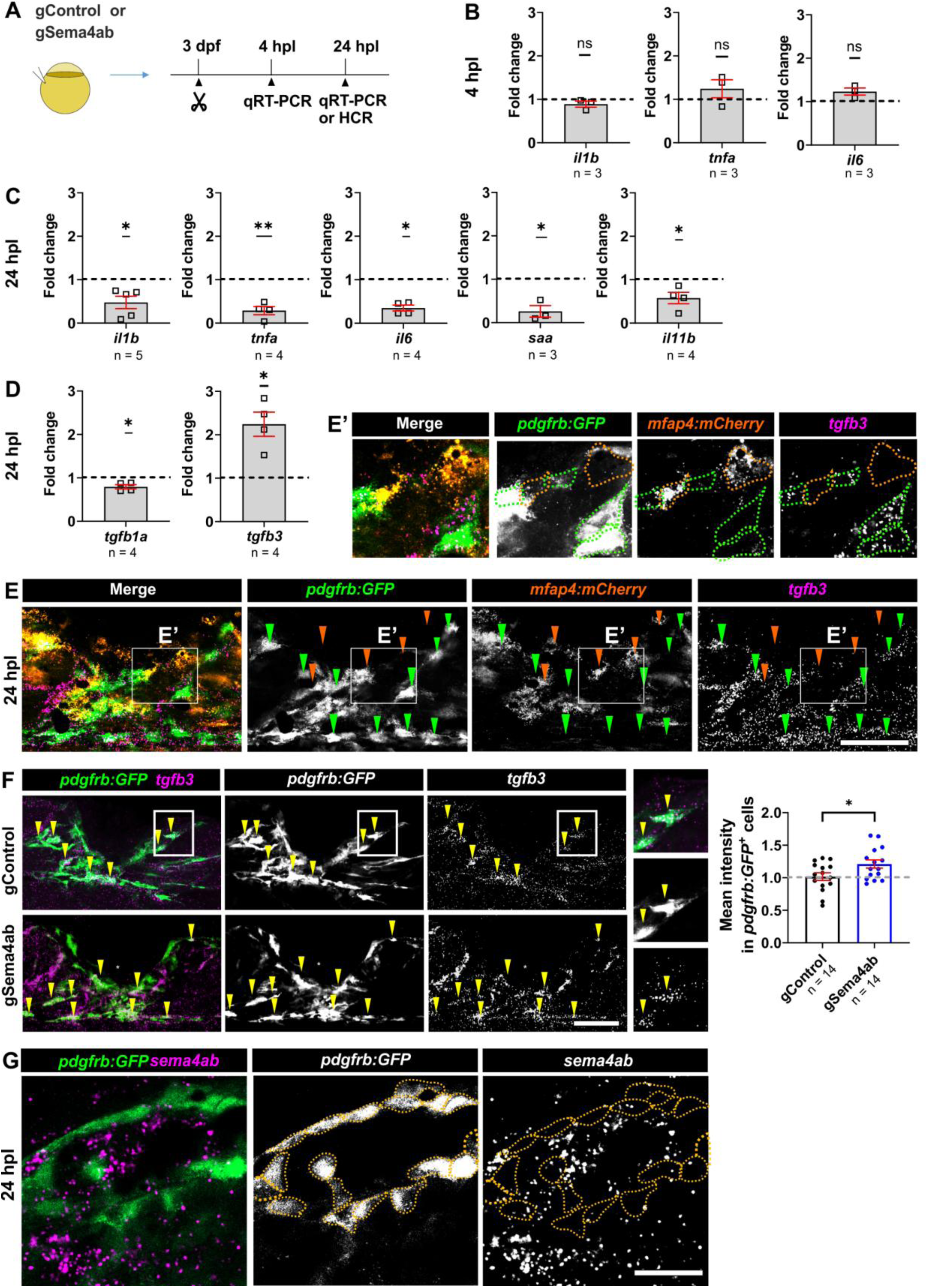
Loss of *sema4ab* function in microglia changes the cytokine profile of the lesion site and increases *tgfb3* expression fibroblasts. **(A)** A schematic indicating the experimental design for cytokine quantification by qRT-PCR is shown. **(B)** qRT-PCR analyses of major pro-inflammatory cytokines in *sema4ab* somatic mutants show no differences to injured control gRNA-injected larvae at 4 hpl (One-sample t-test; *il1b*: p = 0.2526; *tnfa*: p = 0.3623; *il6*: p = 0.1028)**. (C)** At 24 hpl, *sema4ab* somatic mutants show decreased expression levels of pro-inflammatory cytokines and markers (One-sample t-test*; il1b*: −2.12 fold-change, p = 0.0212; *tnfa*: −3.58 fold-change, p = 0.0047; *il6*: −2.94 fold-change, p = 0.0024; *saa*: −3.84 fold-change, p = 0.0311; *il11b*: −1.73 fold-change, p = 0.0495). **(D)** While the anti-inflammatory cytokine *tgfb1a* shows a slight decrease (−1.26 fold-change, One-sample t-test; p = 0.0251), *tgfb3* is substantially increased in expression in *sema4ab* somatic mutants (+2.24 fold-change, p = 0.0210). (**E, E’)** HCR-FISH against *tgfb3* in the *pdgfrb:GFP* x *mfap4:mCherry* double transgenic line shows expression of *tgfb3* only in *pdgfrb:GFP*+ cells (green arrowheads and green dotted circles) and not in *mfap4:mCherry*+ cells (orange arrowheads and orange dotted circles). **(F)** HCR-FISH at 24 hpl shows *tgfb3* expression in fibroblast-like cells (*pdgfrb:GFP*^+^; arrowheads). The mean intensity of the *tgfb3* signal in *pdgfrb:GFP*^+^ cells shows an increase in *sema4ab* haCR-injected larvae at 24 hpl (+1.22-fold change, p = 0.0322). **(G)** HCR-FISH against *sema4ab* in the *pdgfrb:GFP* transgenic line shows hardly any cellular expression of *sema4ab* in *pdgfrb:GFP*+ cells (orange circles). Each dot for qRT-PCR represents a pool of 50 larvae. Error bars show SEM. Scale bars: 50 µm.

Interestingly, after co-disruption of *plxnb1a/b,* cytokine expression levels of *il1b, tnfa, il6 and tgfb3* remained unchanged in these animals in qRT-PCR (Suppl. Fig. 8D, E). Hence, *plxnb1a/b* do not mediate the *sema4ab*-dependent changes in the inflammatory milieu, supporting that their action on neurogenesis could be due to ligand-receptor interactions in ERGs and not by changing the inflammatory milieu of the lesion.

### Fibroblasts increase *tgfb3* expression in the absence of *sema4ab* function in immune cells

Next, we analysed the cell type(s) expressing *tgfb3*, since scRNA-seq predicted the cytokine not to be strongly expressed in immune cells. In HCR-FISH, the labelling intensity of *tgfb3* mRNA was generally increased in *sema4ab* somatic mutants compared to control gRNA injected controls, in agreement with qRT-PCR results (Suppl. Fig. 9A). Remarkably, expression of *tgfb3* in fibroblasts (*pdgfrb*+), but not in BDMs or microglia was observed in scRNA-seq (see Fig. 2F). In agreement with scRNA-seq, HCR-FISH co-localisation studies indicated that the *tgfb3* signal was not detectable in BDM/microglia cells (*mfap4*:*mCherry*+). Instead, the signal was present mostly in fibroblasts (*pdgfrb:GFP*+; Fig. 8E, E’) and increased by 20% in the absence of *sema4ab* function (Fig. 8F). Conversely, *sema4ab* was hardly detectable in fibroblasts in the lesion by HCR-FISH (Fig. 8G). This suggests a scenario in which expression of *sema4ab* in microglia influences the lesion site environment in such a way that *tgfb3* production by fibroblasts is reduced.

Indeed, live observation of BDMs/microglia (*mfap4:mCherry*) and fibroblasts *(pdgfrb:GFP)* from 10 to 25 hpl indicated close physical interactions between these cell types in the injury site, where *mfap4:mCherry+* cells move along elongated stationary fibroblasts (Suppl. Fig. 9B, C and supplemental movies 1 and 2). No obvious differences in these cell behaviours were detectable between control gRNA- and *sema4ab* haCR-injected larvae. These data suggest that BDMs/microglia are in close contact with fibroblasts and may modify the microenvironment, such that *tgfb3* expression in fibroblasts is attenuated in a *sema4ab*-dependent manner.

### Fibroblasts promote regenerative neurogenesis via *tgfb3*

The observed increase of *tgfb3* expression levels in fibroblasts after *sema4ab* disruption could affect neurogenesis (Lee et al., 2020). To test the role of *tgfb3* in regenerative neurogenesis in the zebrafish spinal cord, we effectively disrupted the gene in somatic mutants, as shown previously (Keatinge *et al*., 2021). We verified effective gene disruption here by demonstrating resistance of the target site of the haCR to restriction enzyme digestion in RFLP and by showing a 70 % reduced *tgfb3* mRNA expression in qRT-PCR (Suppl. Fig. 9D). In addition, we used a temporally conditional way to reduce Tgfb signalling by pharmacologically blocking Tgfb receptors with the small molecule inhibitor SB431542 (Sun *et al*, 2006). Somatic mutants for *tgfb3* showed a 45 % reduction in the number of newly generated motor neurons (EdU+/*mnx1:GFP*+) after spinal lesion, compared to control gRNA-injected lesioned larvae at 48 hpl (Fig. 9A). Similarly, adding SB431542 to lesioned larvae reduced the number of newly generated motor neurons by 45 %, compared to DMSO treated controls, at 48 hpl (Fig. 9B). These observations support a promoting role of *tgfb3* for regenerative neurogenesis.

**Fig. 9.**
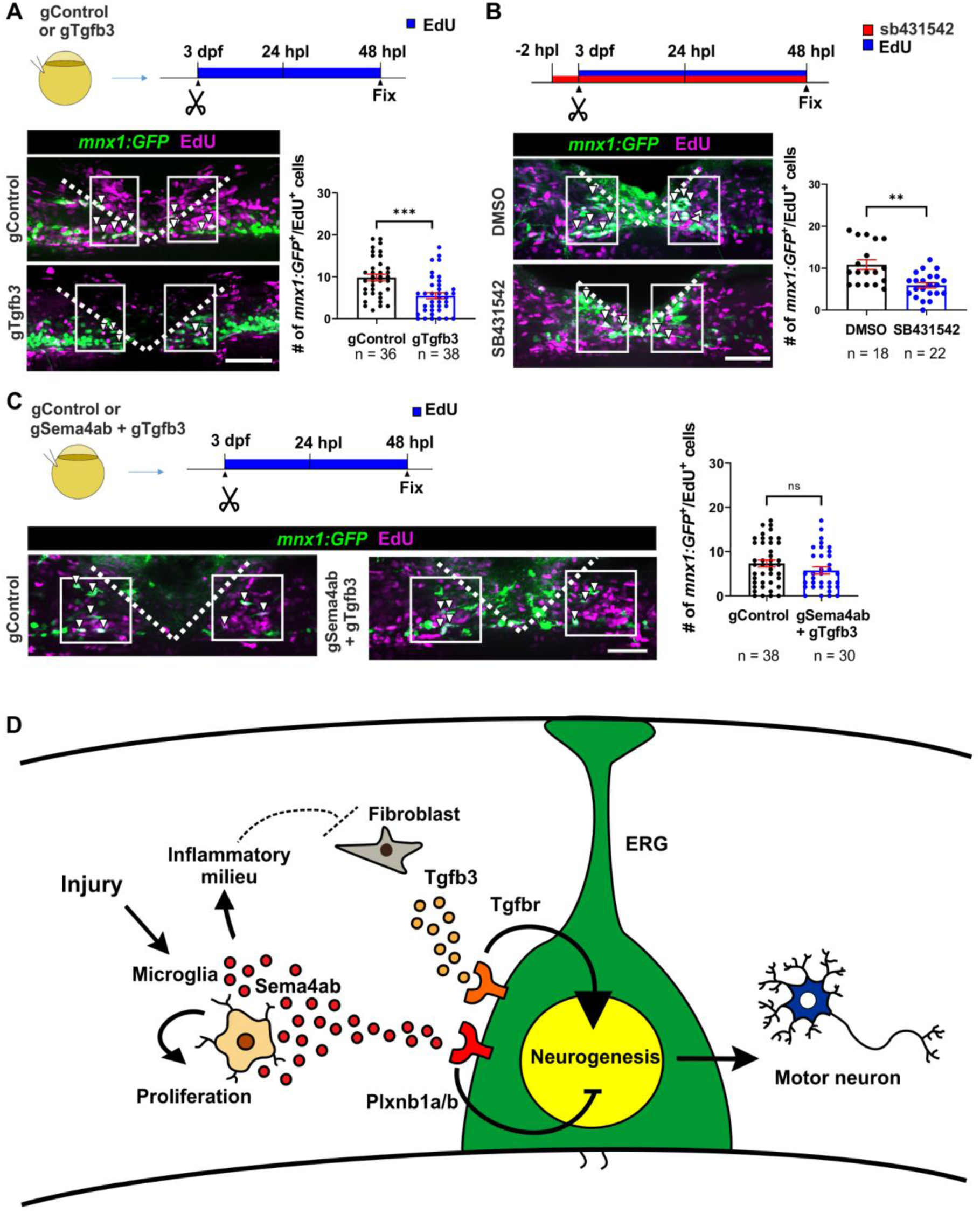
Fibroblasts promote regenerative neurogenesis via *tgfb3*. **(A)** Somatic mutants for *tgfb3* show a reduction in the number of newly generated motor neurons (gControl: 9.8 cells per larva ± 0.8; gTgfb3: 5.5 cells per larva ± 0.7; p = 0.0001). **(B)** SB431542 treatment reduces regenerative neurogenesis (DMSO: 10.8 cells per larva ± 1.15; SB431542: 5.9 cells per larva ± 0.63; p = 0.0011). **(C)** Double haCR injection against *sema4ab* and *tgfb3* restores control levels of regenerative neurogenesis (gControl: 8.1 cells per larva ± 0.7; gSema4ab+gTgfb3: 6.6 cells per larva ± 0.8; p = 0.1523). **(D)** A graphical summary of proposed interactions of *sema4ab* during spinal cord regeneration is shown. After spinal injury, microglia likely control neurogenesis directly by signalling through Sema4ab and Plxnb1a/b on ERGs, and indirectly by changing the injury site environment, which leads to lower expression of *tgfb3*, a positive regulator of regenerative neurogenesis, in fibroblasts. Additionally, Sema4ab controls microglia number by promoting cell proliferation. Error bars show SEM. Dotted lines show the injury site. Scale bars: 50 µm.

### Disruption of *tgfb3* abolishes the effect of *sema4ab* disruption on lesion-induced neurogenesis

To determine whether control of *tgfb3* expression by Sema4ab signalling could explain at least in part the role of *sema4ab* in attenuating regenerative neurogenesis, we disrupted both genes simultaneously by co-injecting *sema4ab* and *tgfb3* haCRs. Disruption of both genes was effective, as indicated by internal RFLP controls for both targets. In these larvae, lesion-induced generation of motor neurons (EdU+/*mnx1:GFP+*) was comparable to those in lesioned control gRNA-injected larvae (Fig. 9C), fully abolishing the enhancing effect of *sema4ab* disruption on neurogenesis. These data show that *tgfb3* is a strong neurogenesis-promoting signal in fibroblasts and that it is controlled via *sema4ab* expression in microglia.

Interestingly, the reduction of neurogenesis when *sema4ab* and *tgfb3* were co-disrupted was stronger than expected, given the potential contribution of a *sema4ab* to *plxn1b1a/b* direct signalling axis, which would still be disrupted in the absence of *sema4ab*. A likely reason for that is that haCRs almost completely disrupt gene function and cannot be titrated to match expression levels of *tgfb3* present in control lesioned animals. Overall, these observations suggest that control of *tgfb3* expression in fibroblasts is a major mechanism by which *sema4ab+* microglia attenuate lesion-induced neurogenesis.

### Potential *sema4ab* signalling in adult zebrafish and mouse spinal cord **injury**

To determine whether cell type-specific expression of *sema4ab, plexins* and *tgfb3* is conserved across ontogeny and phylogeny, we analysed expression of relevant genes in published scRNA-seq profiles of the lesioned spinal cord of adult zebrafish (Cigliola *et al*., 2023) and adult mice (Xue *et al*, 2024).

In adult zebrafish, we found expression of *plxnb1a* and *plxnb1b* in clusters of *her4.1*+ ERGs. In scRNA-seq data, *sema4ab* reads were present in microglia (*p2ry12+*)*. tgfb3* reads were present in fibroblasts (*pdgfrb+*), but not in immune cells (Suppl. Fig. 10A,B). We confirmed expression of *plxnb1a and plxnb1b* this by HCR-FISH in cross sections of adult spinal cord, where we found expression in the ventricular layer (Suppl. Fig. 10C).

Similarly, in the sample from the lesioned spinal cord of adult mice, we found selective expression of *Sema4A* in microglia (*P2ry12+*) and *Plxnb1* selectively in potential ventricular progenitors (*Foxj1+*). *Tgfb3* was expressed in fibroblasts (*Pdgfra+*), but not in microglia (Suppl. Fig. 11A,B). This indicated that in the injured adult zebrafish and mouse spinal cords, the relevant genes are expressed in comparable cell types. However, whether expression levels and dynamics are sufficient for similar interactions between cell types as we observe in larval zebrafish to occur in other systems needs to be directly tested.

In summary, we propose a model by which Sema4ab from microglia controls neurogenesis directly through Plxnb1a/b on ERGs and indirectly via control of fibroblast-derived Tgfb3 in successful spinal cord regeneration in zebrafish (Fig. 9D).

## DISCUSSION

Here we elucidate a mechanism by which regenerative neurogenesis is controlled in a regeneration-competent vertebrate. We show that *sema4ab*, mainly expressed by microglia, influences several cell types. We provide evidence for major functions of Sema4ab in preventing ERG progenitor over-proliferation directly via Plxn1a/b receptors on ERGs and indirectly, by inhibiting Tgfb3 signalling from fibroblasts.

Even though mostly regeneration-promoting signals are reported for regenerating systems, regulation of growth is indispensable for appropriate repair. Here we show that expression of *sema4ab* attenuates regenerative neurogenesis. Some other attenuating signals have also been reported. For example, notch signalling is re-activated after spinal injury in adult zebrafish and attenuates neurogenesis there (Dias *et al*, 2012) and *myostatin a* is a secreted factor by spinal ventricular cells that also attenuates regenerative neurogenesis (Saraswathy *et al*., 2022). In a recent functional screen in the regenerating retina of adult zebrafish, 11 out of 100 regeneration-associated genes have been found to promote and 7 genes to attenuate neuronal regeneration (Emmerich *et al*, 2024). Interestingly, when a stroke is induced or Notch signalling is disrupted, limited neurogenesis can be elicited from ependymal cells in the lateral ventricles of mice, but this leads to depletion of the ependymal cells (Carlen *et al*., 2009). Hence, attenuating factors are likely needed to keep the balance between progenitor maintenance and neuronal differentiation in successful regenerative neurogenesis.

Microglia may signal directly to ERGs to attenuate regenerative neurogenesis. Both signal (*sema4ab*) and receptors (*plxnb1a/b*) were expressed with high selectivity in microglia and ERGs, respectively, as indicated by HCR-FISH. However, other cell types may also express low levels of these genes, as suggested by scRNA-seq. Direct signalling from BDMs to ERGs has been shown for Tnfa, but with a promoting effect on neurogenesis (Cavone *et al*., 2021). In mammals, microglia also attenuate regenerative neurogenesis in the subventricular zone and hippocampus, which is related to the microglia’s pro-inflammatory status and may in part explain the poor regenerative outcome there (Matsuda *et al*, 2015; Monje *et al*, 2003; Nath *et al*, 2024).

Sema4ab/Plxn1a/b interactions inhibit proliferation of neural progenitor ERGs, but an additional effect on neuronal differentiation cannot be excluded. Interestingly, in mice, mutating *PlxnB1* and *PlxnB2* together decreased proliferative activity of progenitors, but increased their differentiation in the developing cortex, leading to a net effect of reduced neurogenesis (Daviaud *et al*., 2016). Our re-analysis of mouse scRNA-seq data shows that microglia in a spinal injury site in mice selectively express *Sema4A* and spinal progenitors express *Plxnb1*. Hence, in principle, Sema4A could also contribute to inhibition of neurogenesis in the lesioned spinal cord of mice. A relatively stronger Sema4A signal in mice than in zebrafish could tip the balance to inhibition of neurogenesis after spinal injury.

In addition to potential direct signalling to ERGs, *sema4ab* also maintains the activation state of microglia. KEGG analysis in microglia clusters highlighted reduced activity of important activation pathways, related to bacterial (Toll-like receptors),viral (RIG-I) response, and initiation of the inflammatory response (Alhamdan *et al*., 2024; Almeida-da-Silva *et al*., 2023; Yoneyama *et al*., 2024) to be changed after disruption of *sema4ab*. In addition, the top down-regulated genes, including activation markers such as *grn1* (Purrahman *et al*, 2023), confirm this change in activation state. Finally, we found reduced expression of pro-inflammatory cytokines in microglia. Since the activation state of microglia affects cytokine signalling (Raziyeva *et al*, 2021), it was not surprising to detect significant changes in the abundance of different cytokines, in particular a reduction of pro-inflammatory ones, in the absence of *sema4ab.* In mammals, microglia cells also express Sema4A (Leitner *et al*., 2015; Meda *et al*., 2012), including in a spinal lesion site, as analysed here. In agreement with our findings, injecting exogenous Sema4A into the mouse brain increased activation of microglia there (Chiou *et al*, 2019). Mammalian macrophages also increase *Sema4A* expression when activated by lipopolysaccharides (Meda *et al*., 2012).

Changes in the cytokine milieu are the likely mechanism by which disruption of *sema4ab* leads to global changes in the injury site environment. In scRNA-seq, we could see profound changes in gene expression in many cell types. However, their abundance in the injury site did not change drastically, with the notable exception of a reduction in microglia, as indicated by scRNA-seq and histology. Even debris endocytosis by BDMs/microglia appeared unaltered, although subtle changes, e.g. in terms of cargo, are possible. Hence, scRNA-seq analysis reveals complex signalling interactions in a spinal injury site that depend on microglia activation. This data presents a resource for further analysis of the complex lesion site environment.

As an example, we here demonstrate that *sema4ab* signalling regulates expression of *tgfb3*, a crucial signalling molecule, in fibroblast-like cells in vivo (John et al., 2025). We detect *tgfb3* expression only in fibroblasts by HCR-FISH and scRNA-seq, and *sema4ab* mostly in microglia. Hence, changes of *tgfb3* expression after *sema4ab* disruption are most likely a consequence of the perturbed cytokine environment. Interestingly, our re-analysis of mouse scRNA-seq data shows expression of *Tgfb3* in fibroblasts, but not in microglia, and *Sema4A* in microglia, but not in fibroblasts in a spinal injury site, raising the possibility for similar suppression of *Tgfb3* in fibroblasts in an environment that is conditioned by Sema4A from microglia in a non-regenerating system. The interactions between fibroblasts and immune cells after CNS injury in mammals are highly dynamic, such that in vivo experiments would be needed to determine whether expression levels in mice are adequate to establish such interactions (Ewing-Crystal *et al*., 2025).

How microglia signal to fibroblasts is unclear. It is most likely that fibroblasts react to changed levels of pro-inflammatory cytokines in the injury site. For example, responsiveness of fibroblasts to pro-inflammatory cytokines, such as TNF (Osei *et al*, 2020) and IL-1beta (Pilling *et al*, 2015) is well established and CellChat identified potentially altered *tnfb/tnfrsf1a* receptor/ligand signalling between microglia and fibroblasts in the absence of Sema4ab signalling. However, a number of other potential receptors for Sema4ab (*havcr1, havcr2, nrp1b, plxnd1*) (Alto & Terman, 2017; Rajabinejad *et al*., 2020) could also play a role in the complex signalling interactions of different cell types in the injury site.

Our data indicate that fibroblast-produced Tgfb3 promotes regenerative neurogenesis. Exogenous TGFB3 has been shown to be a mitogen for retinal cells of the developing rat *in vitro* (Anchan & Reh, 1995), such that Tgfb3 could act on progenitor cells directly in the lesioned spinal cord. However, fibroblast derived Tgfb3 also mitigates inflammation in the lesioned spinal cord of larval zebrafish, indicating additional systemic functions of the cytokine (John et al., 2025). In the adult zebrafish retina, Tgfb3 has been shown to increase pSmad levels in Müller glia, the retinal progenitor cells, and inhibit their injury-induced proliferation (Lee *et al*, 2020). However, there is also evidence for pro-regenerative functions of Tgfb3 in zebrafish retina regeneration, perhaps through indirect effects (Conedera *et al*, 2021; Lenkowski *et al*, 2013; Sharma *et al*, 2020). This indicates context-dependent functions for Tgfb3 in neural development and regeneration.

The altered cytokine environment is also likely to contribute to reduced axonal regrowth in the absence of *sema4ab* function. This could happen through modulation of the ECM (Shah *et al*, 1995; Weng *et al*, 2020) or other interactions that need to be elucidated.

We conclude that the microglia activation state is crucial for the outcome of a spinal injury. The balance of pro- and anti-regenerative signals from multiple cell types determines the regenerative outcome. Single cell -omics approaches are well placed to unravel such complexity. Here, we identify microglia-derived *sema4ab* as a potentially evolutionarily conserved signal in the regulation of the cytokine milieu and as an inhibitor of regeneration of neurons and axons. Hence, *Sema4a* may present a potential therapeutic target in non-regenerating mammals.

## MATERIALS AND METHODS

### Animals

All zebrafish lines were raised and kept under standard conditions (Westerfield, 2000). Zebrafish experiments were performed under state of Saxony licenses TVV 36/2021, TVV 45/2018, and holding licenses DD24-5131/364/11, DD24-5131/364/12. For experimental conditions, larvae up to an age of 5 days post-fertilization (dpf) were used of the following lines: wild type (AB); Tg(*mnx1:GFP*)^ml2tg^, abbreviated as *mnx1:GFP* (Flanagan-Steet *et al*., 2005); Tg(*her4.1:EGFP*)^y83Tg^, abbreviated as *her4.1:EGFP*) (Yeo *et al*., 2007); Tg(*olig2:EGFP*), abbreviated as *olig2:EGFP* (Shin *et al*, 2003); Tg(*mpx:GFP*)^uwm1^, abbreviated as *mpx:GFP* (Renshaw *et al*., 2006); Tg(*mpeg1.1:mCherry*)^gl23^, abbreviated as *mpeg:mCherry* (Ellett *et al*., 2011); Tg(*pdgfrb:Gal4ff*)^ncv24^; Tg(*UAS:GFP*), abbreviated as *pdgfrb:GFP* (Ando *et al*, 2016); TgBAC(*il1b:eGFP*)^sh445^, abbreviated as *il1b:EGFP* (Ogryzko et al., 2019); and Tg(*tnfa:EGFP*)^ump5Tg^, abbreviated as *tnfa:EGFP* (Nguyen-Chi *et al*, 2015).

### Spinal cord lesion

Lesions of larvae were performed as previously described (Ohnmacht et al., 2016). Briefly, at 3 dpf, zebrafish larvae were deeply anaesthetized in E3 medium (Nüsslein-Volhard & Dahm, 2002) containing 0,02 % of MS-222 (Sigma). Larvae were then transferred to an agarose plate. After removal of excess water, larvae were placed in lateral positions. A transection of the entire spinal cord was made using a 30G syringe needle at the level of the 15^th^ myotome, without injuring the notochord.

### Generation of somatic mutants

Somatic mutations were generated mainly following a previous protocol (Keatinge et al., 2021). CRISPR gRNAs were selected using the CRISPRscan website (www.crisprscan.org/gene) webtools. gRNAs were chosen based on the CRISPRScan score (preferentially > 60), and on the availability of a suitable restriction enzyme recognition site (www.snapgene.com) that would be destroyed by the gRNA. Additional criteria were to avoid the first exon and to target functional domains. 1 nL of gRNA mix (1 µl of Tracer (5nM, TRACRRNA05N, Sigma-Aldrich), 1 µl of gRNA (20µM, Sigma-Aldrich), 1 µL of phenol red (P0290, Sigma-Aldrich) and 1 µL of Cas9 (20µM, M0386M, New England Biolabs) was injected into one-cell stage embryos. gRNA efficiency was tested by restriction fragment length polymorphism analysis (RFLP) in a proportion of larvae for each experiment as an internal experimental control (Keatinge et al., 2021). Sequences for gRNAs and primers for RFLPs can be found in Suppl. Table 7. All experiments conducted with somatic mutants were carried out with a control injected group with a non-targeting gRNA.

### Generation of sema4ab germline mutant and Tg(mfap4:mCherry-CAAX;myl7:mCerulean) line

The *sema4ab* mutant stable line (*sema4ab*^tud123^) was generated by raising somatic mutants as potential founders, as described (Keatinge et al., 2021). The founder selected to generate the stable line carried a 14 bp insertion in exon 6 that generated a frame shift with a premature stop codon. Sequencing primers were the same as for RFLP.

BDMs/microglia selectively express *mfap4* (Walton et al, 2015). To generate the *Tg(mfap4:mCherry-CAAX;myl7:mCerulean)* line (abbreviated as *mfap4:mCherry)*, pDEST *mfap4:turquoise2* vector purchased from Addgene (#135218) was used to PCR-amplify the *mfap4* promoter sequence with primers designed to introduce 5’ SacI and 3’ AscI. In parallel, a second plasmid backbone containing *ins:nls-mCerulean; cryaa:mCherry* was digested with SacI/AscI and ligated with *mfap4* using SacI/AscI sites *(mfap4:nls-mCerulean; cryaa:mCherry).* Subsequently, mCherry was PCR amplified by using primers designed to introduce 5’ EcoRI and 3’ CAAX box followed by a stop codon and PacI. Then *mfap4:nls-mCerulean; cryaa:mCherry* were digested and ligated using EcoRI/PacI to mCherry-CAAX fragment *(mfap4:mCherry-CAAX; cryaa:mCherry).* Finally, the newly generated *mfap4:mCherry-CAAX; cryaa:mCherry* construct and *ins1.1:ins-sepHLourin; myl7:NLS-mCerulean* constructs were digested with EcoRI/PacI to yield compatible fragments which were ligated to generate the final construct *(mfap4:mCherry-CAAX;myl7:mCerulean).*The entire construct was flanked with I-Sce sites to facilitate transgenesis and was injected in wild type AB eggs. Several founders were selected based on mCerulean (blue) expression in the heart and mCherry (red) expression in BDMs and the founder with Mendelian segregation in the F1 was selected. Primers are described in Suppl. Table 8.

### *In situ* hybridization chain reaction (HCR-FISH)

Larvae were collected in E3 medium and incubated with 0.003 % 1-phenyl 2-thiourea (PTU) at 6 hpf to suppress melanocyte development. Larvae were fixed overnight in 4 % paraformaldehyde (PFA) at 4 °C and then transferred to 100 % Methanol (MeOH) at −20 °C overnight for permeabilization. The next day, larvae were rehydrated in a graded series of MeOH at room temperature (RT) and subsequent washes in phosphate buffered saline with 0.1 % Tween (PBST). Thereafter, larvae were treated with 500 μL proteinase K (Roche) at a concentration of 30 μg/mL for 45 min at RT, followed by post-fixation in 4 % PFA for 20 min RT and washes with PBST (3 x 5 min). Pre-hybridisation was performed using 500 μL of pre-warmed probe hybridisation buffer (Molecular Instruments (Choi et al., 2018)) for 30 min at 37 °C. The probe sets, which were designed based on the NCBI sequence by Molecular Instruments (Choi et al., 2018), were prepared using 2 pmol of each probe set in 500 μL of pre-warmed hybridisation buffer. Then, buffer was replaced with probe sets and incubated for 16 h at 37 °C. On the following day, samples were washed 4 x 15 min with pre-warmed probe washing buffer (Molecular Instruments) at 37 °C followed by 2 x 5 min in 5x sodium chloride sodium citrate with 0.1 % Tween (SSCT) at RT. Larvae were then treated with 500 μL of room temperature equilibrated amplification buffer (Molecular Instruments, LA, USA) for 30 min at RT. Hairpin RNA preparation was performed following the manufacturer’s instructions. Samples were incubated with Hairpin RNAs for 16 h in the dark at RT. Finally, samples were washed for 3 x 30 min in 5 x SSCT at RT and immersed in 70 % glycerol for imaging on a Zeiss LSM980 confocal microscope.

For HCR-FISH on adult animal, fish were killed, perfusion-fixed with 4% para-formaldehyde, and tissue slices of 50 µm thickness were obtained from dissected unlesioned spinal cord using a vibrating blade microtome (VT1200S, Leica, Wetzlar, Germany), as described (Becker *et al*, 1997). HCR was performed right away following the protocol described above for larvae, with the exception that no permeabilization steps were applied (starting directly with hybridization blocking buffer step).

### Combining EdU with motor neuron and ERG labelling

Labelling newly generated motor neurons followed previous protocols (Ohnmacht et al., 2016). Briefly, lesioned and unlesioned larvae were incubated in E3 medium containing 1 % dimethylsulfoxide (DMSO, Sigma-Aldrich, Taufkirchen, Germany) and 1:100 EdU (Thermo-Fisher, Schwerte, Germany) for 48 h. Samples were fixed in 4 % PFA for 3h at RT and permeabilised in 100 % MeoH at −20 °C for at least overnight. Larvae were then rehydrated, tails were dissected and incubated in proteinase K (Roche, Sigma-Aldrich) for 45 min at RT at a concentration of 10 μg/mL and then postfixed at RT in 4 % PFA for 15 min. Then, larvae were incubated for 20 min at RT in 0.5 % DMSO-PBST and incubated for 2 h at RT with the Click-it Chemistry Edu Alexa 647 kit (Thermo-Fisher, Waltham, USA) as described by the manufacturer. Subsequently, tails were washed in PBST and incubated with blocking buffer solution (2 % BSA, 1 % DMSO, 1 % Triton, 10 % Normal donkey serum, in PBST) for 1 h at RT and then with a chicken anti-GFP antibody, (1:200, Abcam, Cambridge, UK) at 4 °C for 72 hours. Secondary antibody (alexa-488 anti-chicken, 1:200, Jackson Immuno) incubation took place overnight at 4 °C. Finally, samples were washed in PBST and transferred to 70 % glycerol in PBST, mounted and imaged. Imaging was performed using a Widefield Axio Observer (Inverted) with ApoTome 2 Unit (Zeiss). Images were analysed using Fiji (Rueden *et al*, 2017; Schindelin *et al*, 2012). Newly generated motor neurons were counted by assessing the number of double-positive cells for GFP and EdU. Cells were counted manually along the Z-stack, ensuring counting individual cells in single optical sections, in 50 µm windows located at both sides of the injury site (100 µm in total). To mitigate bias, experimental conditions were occluded for the analysis. At least two independent experiments were performed. Individual experiments were merged and analysed for statistical significance.

To label proliferating ERGs, we followed the same experimental procedure, imaging and cell counting analysis as for newly generated motor neurons using the ERG reporter line *her4.1:EGFP* instead of the *mnx1:GFP* transgenic line.

### Immunofluorescence

Samples were fixed in 4 % PFA at 24 hpl for 1.5h at room temperature and permeabilised in 100 % methanol at −20 °C for at least overnight. Larvae were then rehydrated, tails were dissected and incubated for 10 min in 100% chilled acetone at −20 °C. Samples were washed and incubated with 500 μL proteinase K (Roche) at a concentration of 12.5 μg/mL for 45 min at room temperature, followed by post-fixation in 4 % PFA for 20 min at room temperature and washes with PBST (3 x 5 min). Tails were then incubated with blocking buffer solution (2 % BSA, 1 % DMSO, 1 % Triton, 10 % Normal donkey serum, in PBST) for 1 h at room temperature and then with the corresponding antibody, (mouse anti-acetylated tubulin, 1:300, Sigma, T6793; rabbit anti-mCherry, 1:200, Invitrogen, PA5-34974; mouse anti-4C4, 1:50, European Collection of Authenticated Cell Cultures, 7.4.C4) at 4 °C for 72 hours. After washings in PBST, incubation with matching secondary antibody (1:200, Jackson Immuno) took place overnight at 4 °C. Finally, samples were washed in PBST and transferred to 70 % glycerol in PBST, mounted and imaged.

### Generation of heat shock plasmid and treatment

To create the pTol-hsp70l:sema4ab-T2a-mcherry plasmid, we modified the pTol-hsp70l:mCherry-T2A-atho1b vector (Ezhkova *et al*, 2023). We PCR-amplified the *sema4ab* coding sequence from genomic cDNA and PCR-amplified T2a-mcherry with specific primers (Suppl. Table 9). Then, we performed a simultaneous multiple fragment ligation in the opened backbone vector. The final construct pTol-hsp70l:sema4ab-T2a:mcherry was verified by sequencing, qRT-PCR for over-expression and mCherry fluorescence after injection.

### Measurement of axonal regrowth

As an indicator for successful axon regrowth, we determined the extent of continuous labelling of axons across the spinal cord injury site, termed axonal bridging. We determined the thickness of the axonal bridge at its narrowest point. Spinal lesion was performed at 3 dpf as described in larvae above. At 2 hpl, larvae were inspected under a fluorescence stereo-microscope and those larvae with incomplete spinal cord transection and those in which the notochord was inadvertently injured (over-lesioned) were excluded from the experiment. At 48 hpl, larvae were fixed and immuno-labelled against acetylated tubulin. Images were taken with a Dragonfly Spinning Disk microscope (Andor/Oxford Instruments, Belfast, UK) with a step size of 1 µm.

The bridge thickness was determined using a custom script in Fiji (Rueden *et al*., 2017; Schindelin *et al*., 2012). Measures were taken at the narrowest point of the axonal bridge, automatically detected by using a maximum projection of the z-stack and a binary segmentation. Measurements of three independent experiments were obtained and merged for statistical analysis.

### Analysis of swimming behaviour

The swimming behaviour was assessed using the ZebraBox (View Point, Lyon, France). As above, under- and over-lesioned larvae were excluded from the experiment at 2 hpl. Subsequently, the larvae were individually arranged in 24-well plates, ensuring representation of all groups on each plate, and kept in the incubator at 28 °C until 48 hpl. Larval movements were automatically tracked during three repetitions of the same protocol. The protocol comprised 1 min of rest, a 1 second vibration stimulus (frequency set at 1000 Hz [target power 100]), followed by 59 seconds of no stimulus. The analysis focused on the time-period of 1 minute encompassing the vibration stimulus (including the 1-second stimulus and the subsequent 59 seconds). For each larva, the mean of their swimming distances over the three repeats was computed. Two independent experiments with an average of 30 larvae per condition were performed and the results were merged for statistical analysis (see below and figure legends).

### Quantitative RT-PCR

Following spinal cord lesion of 3 dpf, RNA extraction was performed at 1 dpl using the RNeasy Mini Kit (Qiagen) according to the manufacturer’s instructions (50 larvae/condition). The injury site was enriched by cutting away rostral and caudal parts from the larvae at positions approximately 3 somites away from the lesion site in either direction. cDNA was synthesized using the iScript™ cDNA Synthesis Kit (BioRad, cat #. 1708890) according to the manufacturer’s instructions. qRT-PCR was performed using the SsoAdvanced Universal SYBR Green Supermix kit (BioRad) according to the manufacturer’s instructions and run in triplicates on a LightCycler 480SW instrument (Roche). For gene-specific primers see Suppl. Table 10. Each condition was normalized to a housekeeping gene and the experimental group was compared to controls by normalizing to the control group. At least three individual experiments were run. For statistical analysis, a One-Sample T-test was used.

### Immune cell counting in whole-mounted larvae

Larvae were lesioned at 3 dpf and fixed for 1 h in 4 % PFA at different time points. Larvae were washed in PBST and transferred to glycerol (70 % in PBST), mounted and imaged immediately using a Widefield Axio Observer (Inverted) with an ApoTome 2 Unit. Images were analysed using Fiji with the experimental condition unknown to the observer. A window of 200 µm width and 75 µm height was placed in the centre of the injury site of individual images, right above the notochord. Cells were counted manually inside the window along the Z-stack with the experimental condition occluded. At least two individual experiments were carried out and merged.

### Acridine Orange labelling

Larvae of the *mpeg:mCherry* line were incubated after a lesion in 1 µg/ml of Acridine Orange (stock solution 10 mg/ml in water) in E3 for 24 h. Then, fish were washed for 3 x 10 min in E3. Larvae were then mounted in 0.5 % agarose in E3 for live imaging on the Dragonfly Spinning Disk microscope (Andor). To estimate total debris, GFP mean intensity was analysed using a 250x100 µm counting window placed in the centre of the injury site. To determine the number of BDMs/microglia that ingested debris, cells positive for both mCherry and GFP were counted in the injury site using same window as before.

### Fluorescence intensity/area measurements

Images were analysed with Fiji ImageJ (https://imagej.net/ij/). The region of interest was defined as a rectangle of 400 x 100 µm, centred on the injury site and located right above the notochord. Z-stacks were collapsed in sum intensity projections and then background subtraction was performed before analysing the mean intensity with a custom Fiji macro to automatize the process (Rueden *et al*., 2017; Schindelin *et al*., 2012). Mean intensity was represented by normalizing each value to the control average value.

### Drug treatment of larvae

SB431542 (Fisher Scientific) was dissolved in DMSO to a stock concentration of 10 mM. The working concentration was 50 µM prepared by a dilution of the stock solution in E3. Larvae were pre-treated for 2 h before injury and then incubated for 48 hpl. Control larvae were treated with DMSO at the same concentration.

### Cell dissociation and fluorescence-activated sorting

50 larvae per sample were used. Lesion-site tissue was enriched by dissecting 3 somites rostral and caudal of the injury site and collecting them in cold 1 x PBS. The tissue was dissociated using the Miltenyi Neural Tissue Dissociation Kit (P) with addition of collagenase. In brief, we removed excess 1x PBS and added 950 µL Buffer X. 6 µL of 1/1000 diluted β-mercaptoethanol, 50 µL Enzyme P and 8 µL collagenase (50 mg/mL) were added. The tissue was pipetted 10 times with 1 mL pipette tips and let it rotate at RT (∼22-23 °C) for approximately 1 min. In the meantime, 10 µL of Buffer Y and 10 µL of Enzyme A were mixed and added this to the dissociation mix. Tissue was triturated slowly using fire polished small-orifice glass Pasteur pipette. Each trituration was performed 10 times (6 seconds each) for 1 min and we let the sample rotate for 1 min. This step was repeated until complete dissociation was achieved. The reaction was stopped after ∼14 mins by adding 1 mL of DMEM + FBS (1 %) + Protease Inhibitor (1 x) mix (DFP), followed by centrifugation at + 4 °C at 500 g for 5 mins. The supernatant was removed and the pellet was resuspended in 1 mL of HBSS (Mg2+/Ca+) + Protease Inhibitor (1x) and then passed through a 40 µm filter equilibrated with 200 µL of DFP. Cells were recentrifuged at +4 °C at 500 g for 5 mins. The pellet was resuspended in 300 µL of DFP. Just before cell sorting, 0.5 µL of propidium iodide (100 µg/mL) was added and mixed well by pipetting. Viable, propidium iodide-negative cells were sorted using FACS Aria (BD Biosciences) using a 100 µm nozzle at flow rate 1. 100,000 cells per sample were sorted into BSA coated 1.5 mL Eppendorf tubes with 500 µL of a 0.4 % BSA solution in 1x PBS. The sorted cells volume was around 200 µL and the final concentration of BSA was 0.28 %. One control gRNA and two *sema4ab* gRNA samples were processed by the Dresden Concept Genome Center for cell encapsulation using the 10X Genomic V3.

### Droplet encapsulation and single-cell RNA sequencing

Cells were pelleted at 500 g for 5 minutes at 4°C, leaving 20 µL of suspension. Cell viability was assessed with trypan blue, and samples with > 80 % viability were prepared for encapsulation. Single-cell RNA sequencing followed the 10x Genomics (Chromium Single-cell Next GEM 3’ Library Kit V3.1) workflow (Zheng *et al*, 2017). In brief, cells were mixed with Reverse Transcription Reagent and loaded onto a Chromium Single Cell G Chip, targeting 10,000 cells per sample. GEM generation, barcoding, reverse transcription, cDNA amplification, fragmentation, and purification were performed according to the manufacturer’s protocol. Libraries were prepared using 25 % of cDNA, quantified, and sequenced on an Illumina NovaSeq 6000 in 100 bp paired-end mode.

### Alignment to zebrafish genome

The raw sequencing data were processed with the ‘count’ command of the Cell Ranger software (v7.1) provided by 10x Genomics. To build the reference, the zebrafish genome (GRCz11) and gene annotation (Ensembl 109), were downloaded from Ensembl (https://www.ensembl.org/). The ‘count’ command was used to generate count matrices that were further used by Seurat V5 for downstream analyses.

### Quality Control and Doublet Removal

For data analyses, cells were kept based on the following conditions: (i) nFeature_RNA >= 200 (number of genes detected per cell), 500 < nCount_RNA < 50000 (number of total reads detected per cell), (ii) percent.mt > 20 % (mitochondrial RNA percentage), (iii) nCount_RNA/nFeature_RNA > 20-fold (an indication of possible doublets), (iv) genes found in fewer than 3 cells were removed from the analyses. Additionally, doublets were determined using scDblFinder V1.18.0 (Germain *et al*, 2021). However, rare cells expressing markers for cell types distinct from the clusters they were allocated to remained and were possible doublets. Cells were removed, if cells other than neurons expressed *elavl3* or *elavl4*, if cells other than red blood cells expressed *hbbe1.3/hbbe2*, if cells other than muscle, fibroblast-like cells or keratinocytes expressed *col12a1a*, *col12a1b*, or if cells other than muscle or fibroblast-like cells expressed *fgl1,* or *vwde*. Using this method an additional ∼5% of cells per sample were removed.

### Seurat Data Analyses

After removing low-quality cells and doublets, we utilized Seurat V5 for data integration using CCAIntegration method and followed its standard pipeline. Briefly, a Seurat object was created and the data were normalized. The top 2000 variable genes were identified, and the data were scaled using all genes, with nCount_RNA regressed out. Clustering was performed at resolution 1, yielding 44 distinct clusters (numbered as 0-43).

### Finding Marker Genes and annotation of clusters

To identify marker genes (the genes which are specific to a cluster compared to the remaining all other cell types), the FindAllMarkers function from Seurat, using the default options was used. The “Wilcoxon rank sum test”, and p_val_adj (p-value adjusted) < 0.1 and avg_log2FC >= 0.25 (average log2 fold change) was used for selecting significantly expressed marker genes. Additionally, genes were selected using pct.1 (percentage in the cluster) > 0.2 and ordered, based on their agvg_log2FC in descending order for each cell cluster. Based on the literature, known cell type marker genes were selected from the top ranked genes and used to annotate the main cell types accordingly.

The main cell types were annotated using the following marker genes and the top-ranked genes for each cluster (Cavone *et al*., 2021; Chen *et al*, 2023; Farrell *et al*, 2018): Ependymoradial Glia (ERG; *fabp7a, sox2*), Enteric Nervous System (ENS; phox2aa, phox2b), Fibroblast-Like Cells (FBL; 3 main clusters, *col12a1a*, *col12a1b*, *pdgfrb*+), Keratinocytes (KC; *pfn1, krt4*), Peridermis (PD; *krt5, krt91, icn2)*, Muscle Cells (MC; m*eox1, mylz3*), Xanthophores (XP; *gch2, aox5)*, Oligodendrocyte (OD; *cldnk, mbpb, mpz*), Kolmer-Agduhr Cells (KA; *npcc, sst1.1*), neutrophils (NP; *lcp1, mpx, lyz*), BDMs/microglia (MM; *lcp1, mfap4, mpeg1.1*), Neurons (NN; *elavl3, elavl4*), Endothelia Cells (EC; *kdrl, cldn5b*), Notochord (NC; s*hha, gas2b*), Red Blood Cells (RBC; *hbbe1.3, hbbe2*) and 5 clusters that remained not annotated (c#28, c#31, c#32, c#34 and c#36).

### Differentially Expressed Gene Analysis (DEG)

The FindMarkers function in Seurat was used to determine the differentially expressed genes using “Wilcoxon rank sum test”. p_val_adj < 0.1 and abs(avg log2FC) >= 0.25 were used for selecting significantly expressed genes.

### Gene Ontology term and KEGG Analyses

For KEGG (Kyoto Encyclopedia of Genes and Genomes) analyses, GOstats (Falcon & Gentleman, 2007), giving all differentially expressed genes (over- or under-represented) using 0.25 log2FC and p_value < 0.05 as input, were used. Enriched terms were selected with a p-value < 0.05.

### Cell-Cell Interaction

CellChat V2.1.0 (Jin *et al*., 2021) was used for cell-cell interaction and a custom cell-cell interaction as described (Cosacak et al. 2019). In brief, the CellChat Database was modified by adding *sema4ab* as ligand and *plxnb1a, plxnb1b, plxnb2a, plxnb2b, plxnd1, havcr1, havcr2, nrp1a and nrp1b* as its receptors. As there was a pathway named SEMA4, the pathway was renamed to “SEMA4AB” for *sema4ab* and to “SEMA4other” for *sema4c/sema4g*. Then, CellChat objects were created for each data set using normalized data from the integrated Seurat object. Finally, the CellChat objects were merged for comparative analyses.

### Cell cycle scoring analysis

Cell cycle scoring analysis was performed using the “Cell-Cycling Scoring” code for Seurat in R (Satija *et al*., 2015) based in a published gene list (Tirosh *et al*, 2016) (loaded in Seurat) and adapted to zebrafish orthologues manually using ZFIN (Sprague *et al*, 2003).

### Time-lapse microscopy

Zebrafish larvae were prepared for time lapse microscopy based on our established protocol (https://ibidi.com/img/cms/support/UP/UP12_Zebrafish_Co_Culture.pdf). Briefly, double transgenic zebrafish larvae (*pdgfrb:GFP*; *mfap4:mCherry*) were lesioned at 3 dpf and anaesthetized with MS-222 (160 mg/L in E3 medium). Low melting point agarose (0.5 %, Biozym Plaque Agarose - Low melting Agarose #840100) was prepared in E3 medium and allowed to cool down to 38 °C before adding MS-222 (final concentration 80 mg/L). Anaesthetized zebrafish larvae were transferred to the agarose and one by one positioned into individual wells of the chambered coverslip. Six larvae were used in total; three control gRNA-injected and three *sema4ab* haCR-injected ones. Microscopy was performed with a Dragonfly Spinning Disk microscope (Andor/Oxford Instruments, Belfast, UK) with a temperature-controlled stage at 28 °C using a 20x objective (20x/0.75 U Plan SApo, Air, DIC, OLYMPUS, Tokyo, Japan). The GFP fluorophore was excited with a 488 nm laser (laser power 20 %) with an exposure time of 180 ms and the mCherry flourophore was excited with a 561 nm laser (laser power 50 %) with an exposure time of 900 ms. We acquired z-stacks of 43 optical sections, with a step size of 1 µm. The interval between single frames of the time-lapse movies was 6 minutes. Time-lapse was performed from 10 hpl until 25 hpl.

### Bias mitigation and statistical analysis

The experimental conditions were occluded before manual counting and analysis and processing was automated where possible (e. g. measurements of axonal regrowth and intensity measurements). For manual counts of EdU+ cells, pilot experiments were performed by two independent experimenters, both showing comparable effect sizes. Experiments were performed in duplicate, if not indicated differently.

Statistical methods to analyse scRNA-seq data are described above. For all other experiments, quantitative data was tested for normality using Shapiro-Wilk’s W-test. Then, parametric (unpaired Student’s t-test or one-way ANOVA) and non-parametric (Mann-Whitney U test or Kruskal-Wallis test) were applied as appropriate. Error bars in figures indicate standard error of the mean (SEM), p-values were represented as asterisks in the figures (*p < 0.05; **p < 0.01; *** p < 0.001) and exact p-values are given in figure legends. To generate graphs and for statistical analysis we used GraphPad Prism 8 (GraphPad Software, Boston, USA).

### Figure preparation

Images were adjusted for brightness, contrast and intensity. Drawings were made using CorelDraw X8 (Corel/Alludo, Ottawa, Canada). Figure plates were prepared using CorelDraw X8 (Corel/Alludo, Ottawa, Canada) and Adobe Photoshop 2024 (Adobe, CA, USA).

### Data availability

The scRNA-seq raw data of zebrafish larvae generated in this study can be accessed at NCBI GEO accession no. xx. Additional datasets used in this are publicly accessible at NCBI GEO accession numbers; GSE213435 (adult zebrafish), GSE218584 (adult mouse). The R-source codes for single-cell analyses are available on GitHub; https://github.com/micosacak/scRNA_seq_zebrafish_sema4ab.

## ACKNOWLEDGEMENTS

We thank Dr. Stefan Hans for reagents; Drs Hella Hartmann and Ruth Hans for imaging advice; Dr. Daniel Wehner for critical reading; and Dr. Judith Konantz, Marika Fischer, and Silvio Kunadt for fish care. This work was supported by the Light Microscopy Facility, the DRESDEN-concept Genome Center, the Flow Cytometry Facility, and the Zebrafish Facility, all core facilities of the Center for Molecular and Cellular Bioengeneering (CMCB) at the Technische Universität (TU) Dresden. Funding was provided by the BBSRC (BB/T015594/1 to TB, CGB), an Alexander von Humboldt Stiftung Professorship award (to CGB), and TU Dresden core funding (to CGB).

## AUTHOR CONTRIBUTIONS (CRediT nomenclature)

ADS: conceptualization, methodology, validation, formal analysis, investigation, writing - original draft, writing - review and editing, visualisation, supervision; MIC: methodology, validation, formal analysis, investigation, writing - review and editing; KH: methodology, validation, formal analysis, investigation, visualization; FK: software; methodology; AMO: investigation; MW: investigation; ÖC: investigation, DZ: investigation; JA: investigation, AB: investigation; formal analysis; AH: resources; NN: resources; CGB: conceptualization, writing - original draft, writing - review and editing, visualisation, supervision, funding acquisition; TB: conceptualization, writing - original draft, writing - review and editing, visualisation, supervision, funding acquisition.

## SUPPLEMENTARY FIGURES

**Suppl. Fig. 1.**
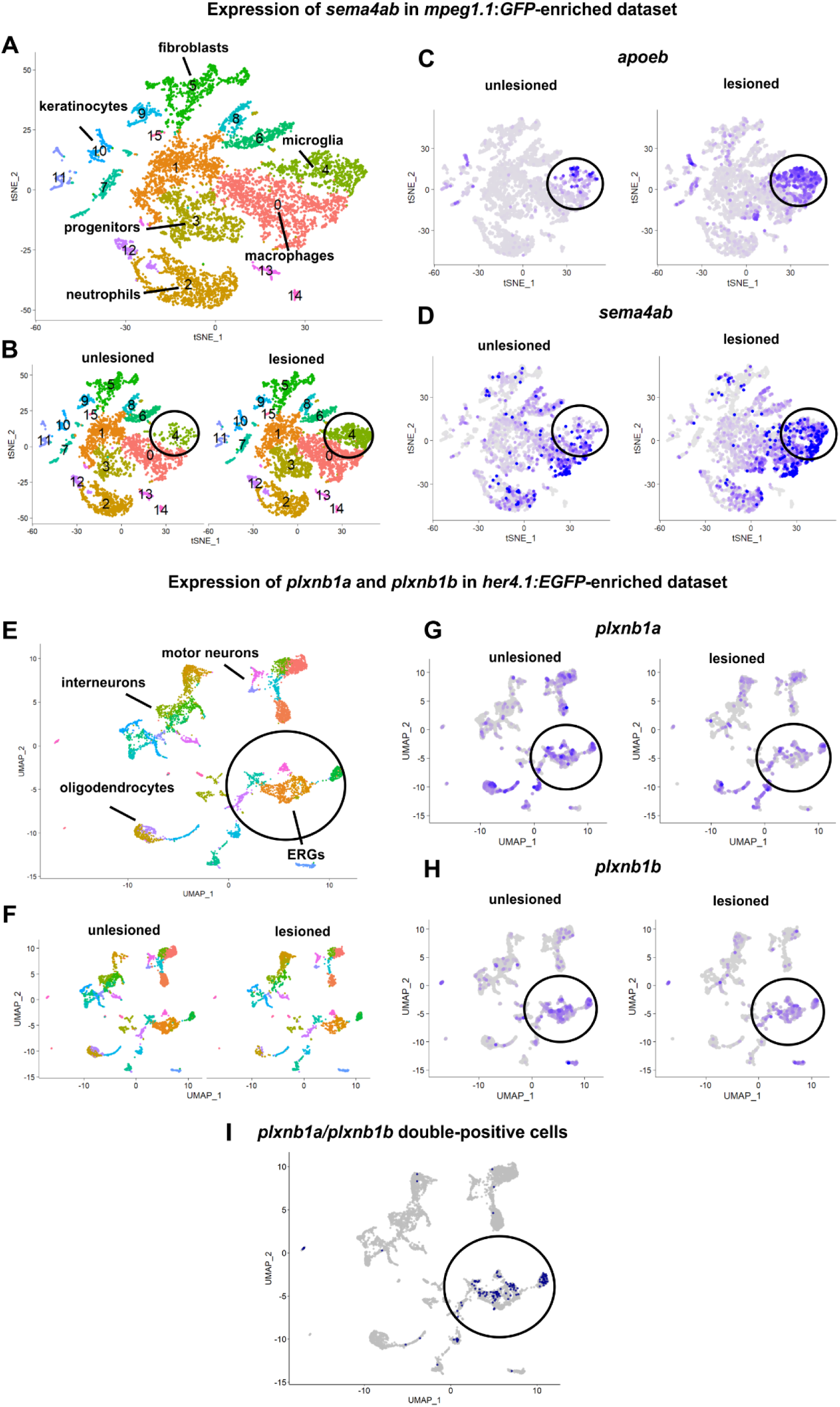
Potential interaction between *sema4ab* in microglia and plexin receptors in ERGs revealed in cell type-enriched scRNA-seq datasets. **(A)** UMAP showing clusters in an *mpeg1.1:GFP*-enriched scRNA-seq. **(B)** UMAPs comparing unlesioned and lesioned conditions are shown. Note an increase of the microglia cluster after injury (circle). **(C,D)** Feature plots comparing expression of *apoeb* (C) and *sema4ab* (D) between unlesioned and lesioned larvae, indicating increased presence of both after injury (circled). **(E**) UMAP showing clusters in a *her4.1:EGFP*-enriched scRNA-seq. Circle indicates ERGs. **(F)** UMAPs indicate presence of ERGs in unlesioned and lesioned conditions. **(G-I)** Feature plots indicating expression of *plxnb1a* (G) and *plxnb1b* (H) in ERGs of unlesioned and lesioned larvae (circled). Plotting only cells co-expressing receptors shows strong enrichment in ERGs (I, circled). Data are described in Cavone et al., Dev Cell 2021 Jun 7;56(11):1617-1630.e6.

**Suppl. Fig. 2.**
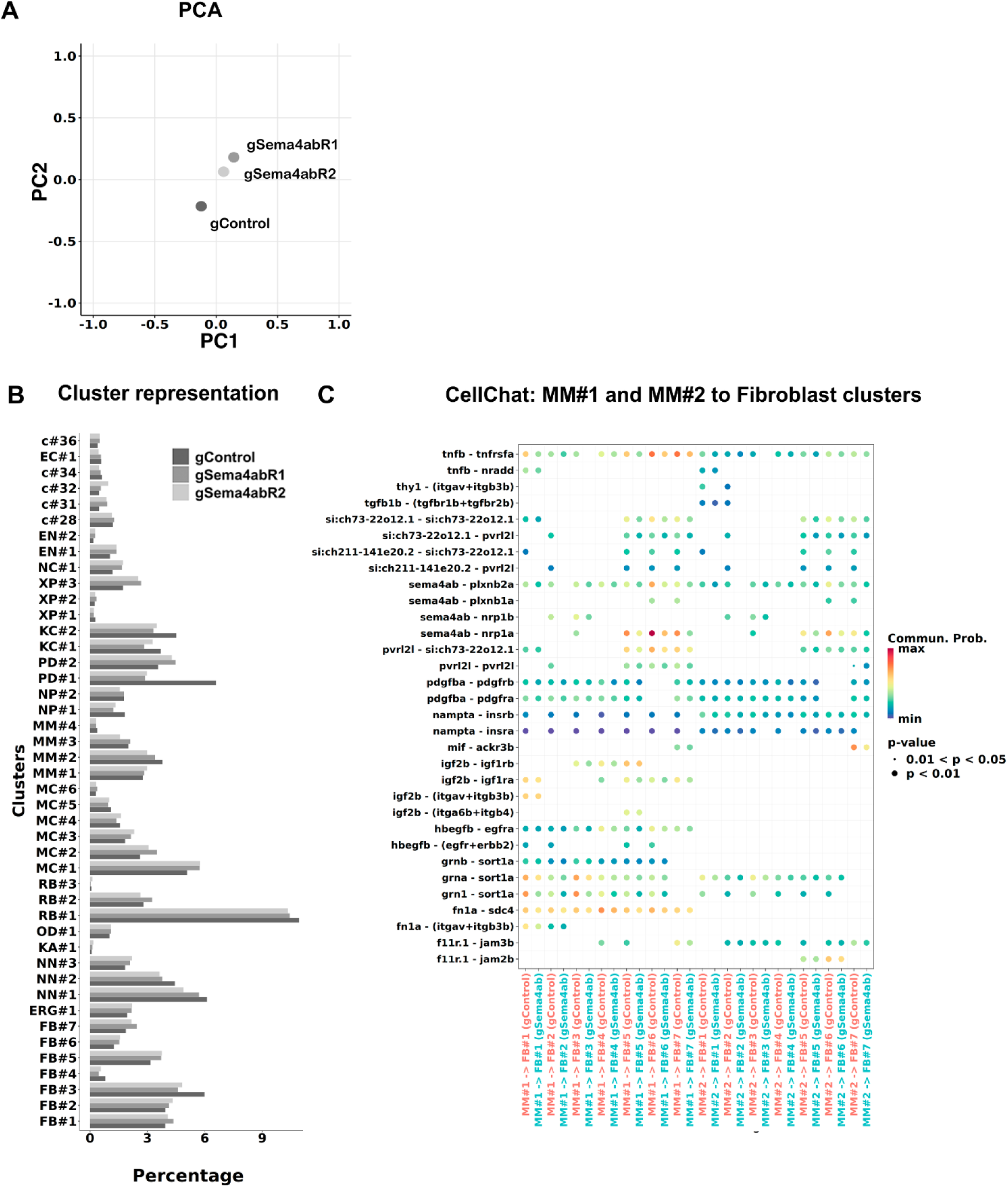
Cell representation and potential interactions in scRNA-seq after *sema4ab* disruption. **(A)** Principal component analysis (PC1 and PC2) shows that gSema4ab replicates (R1 and R2) are more closely related to each other than to gControl. **(B)** A graph indicates similar percentages of cells per cluster across scRNA-seq samples. **(C)** A dot plot illustrates the communication probabilities of all ligand-receptor interactions identified by CellChat between microglia clusters MM#1 and MM#2 and fibroblast clusters. No dot indicates no interaction detected.

**Suppl. Fig. 3.**
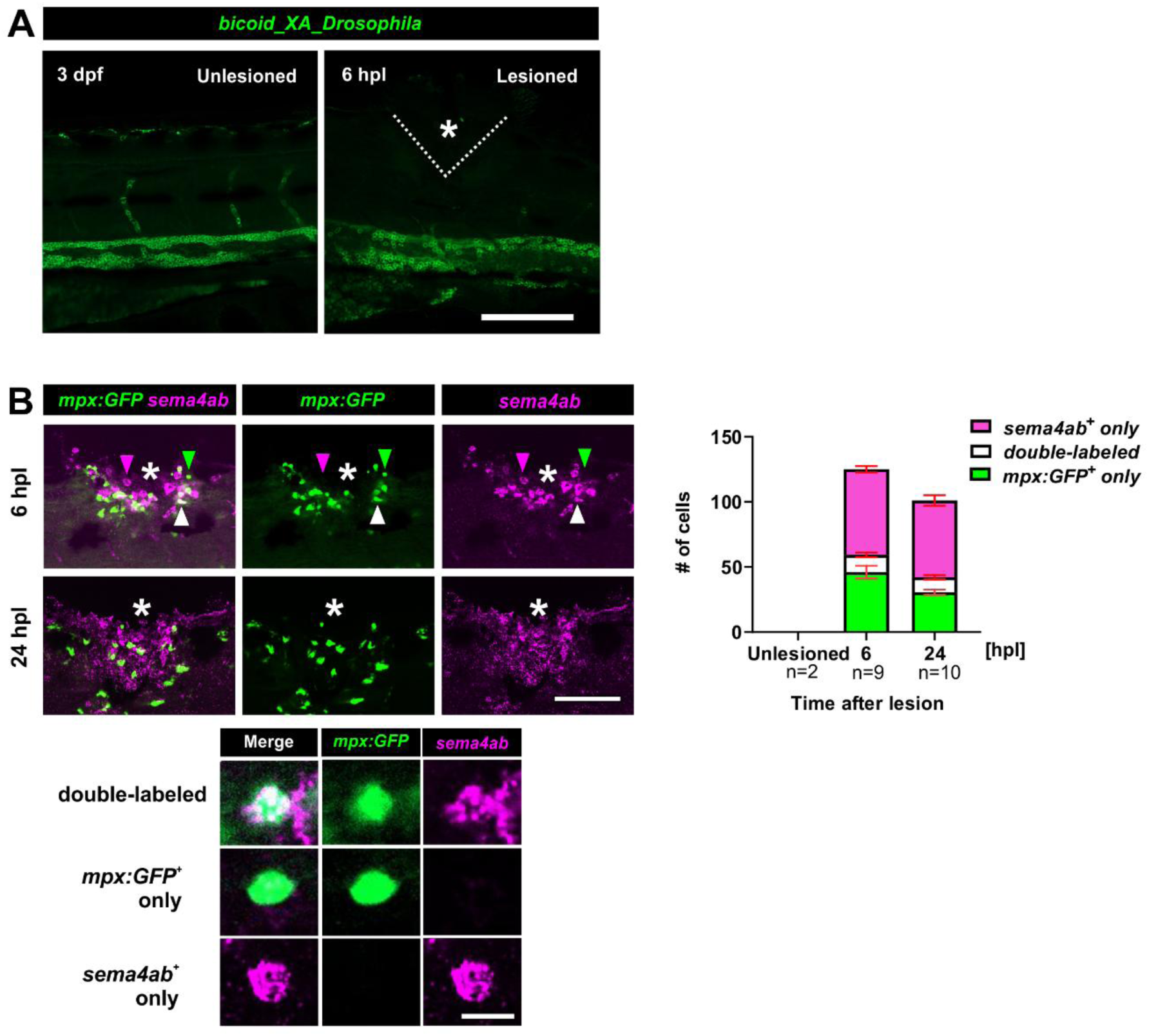
HCR-FISH method control and detection of *sema4ab* expression in neutrophils. **(A)** HCR-FISH for the negative control gene (*Drosophila* BicoidXA) shows non-specific labelling in some blood vessels, but not in the spinal cord in uninjured and injured larvae. **(B)** Multiplexed HCR-FISH at 6 and 24 hpl shows that some *sema4ab+* cells express *mpx:GFP* (white arrows: double-positive cells; green arrows: *mpx:GFP+* only; magenta arrows: *sema4ab*+ only). Scale bars: 100 µm (A,B), 10 µm (inset).

**Suppl. Fig. 4.**
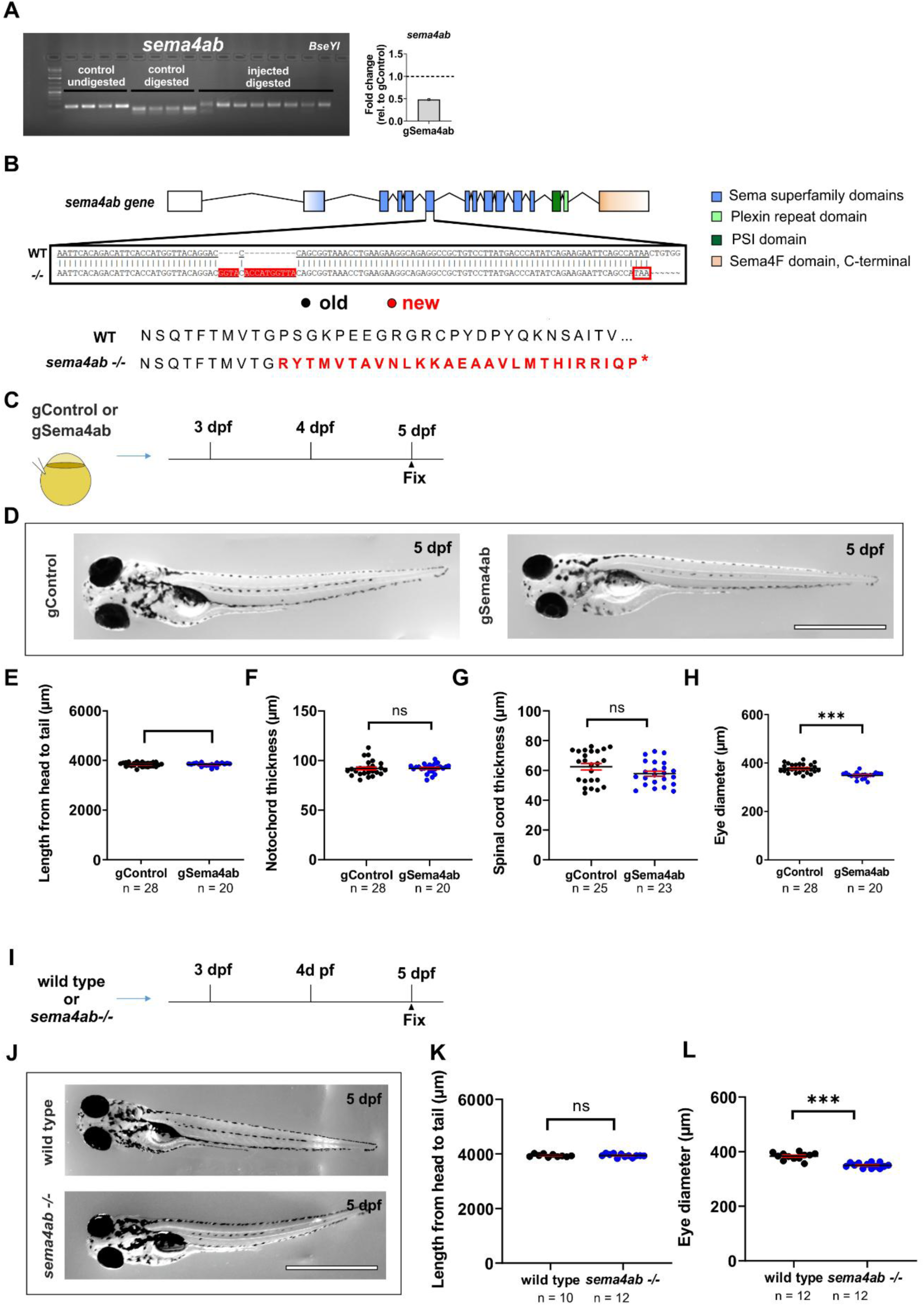
Efficient disruption of *sema4ab* does not lead to large changes of larval growth. **(A)** RFLP and qPCR showing gRNA injection efficiency for *sema4ab*. For RFLP each lane represents one larva with and without digestion with the indicated restriction enzymes and with and without targeting these sites with gRNAs as indicated. Targeting the recognition sites with haCR gRNAs leads to efficient somatic mutation, as indicated by the almost complete resistance to digestion. Note a **∼**50% reduction in mRNA abundance detected by qRT-PCR. **(B)** A schematic of the *sema4ab* gene structure and position of the mutation in the *sema4ab* germline mutant is shown. Note the insertion of 14 bp in exon 6 (sequence underlaid in red) and the frameshift that predicts a stop codon (red frame and asterisk). **(C)** A schematic indicating the experimental design for D-H is shown. **(D)** Photomicrographs of gControl and gSema4ab larvae are shown at 5 dpf. **(D-H)** Quantifications show no differences in body length (E: gControl: 3836 µm ± 15.65; gSema4ab: 3838 µm ± 15.72; t-test: p = 0.9917), notochord thickness (F: gControl: 92.09 µm ± 1.527; gSema4ab: 92.20 µm ± 0.9314; Mann-Whitney U-test: p = 0.3423) or spinal cord thickness (G: gControl: 62.49 µm ± 2.215; gSema4ab: 57.83 µm ± 1.677; Mann-Whitney U-test: p = 0.1134) between gControl and gSema4ab larvae. Eye diameter is slightly reduced in gSema4ab larvae (H: gControl: 378.2 µm ± 3.320; gSema4ab: 349.3 µm ± 2.784; t-test: p < 0.0001). **(I)** A schematic indicating the experimental design for I-K is shown. **(J)** Photomicrographs show wild type and germline mutant larvae for *sema4ab* at 5 dpf. **(K)** No difference was observed in body length of wild type and mutant larvae (wild type: 3941 µm ± 15.95; *sema4ab -/-*: 3940 µm ± 17.50; t-test: p = 0.6855). **(L)** Eye diameter of mutants is slightly decreased (wild type: 383.3 µm ± 3.788; *sema4ab -/-*: 349.9 µm ± 2.56; t-test: p < 0.0001). Error bars show SEM. Scale bars = 1 mm.

**Suppl. Fig. 5.**
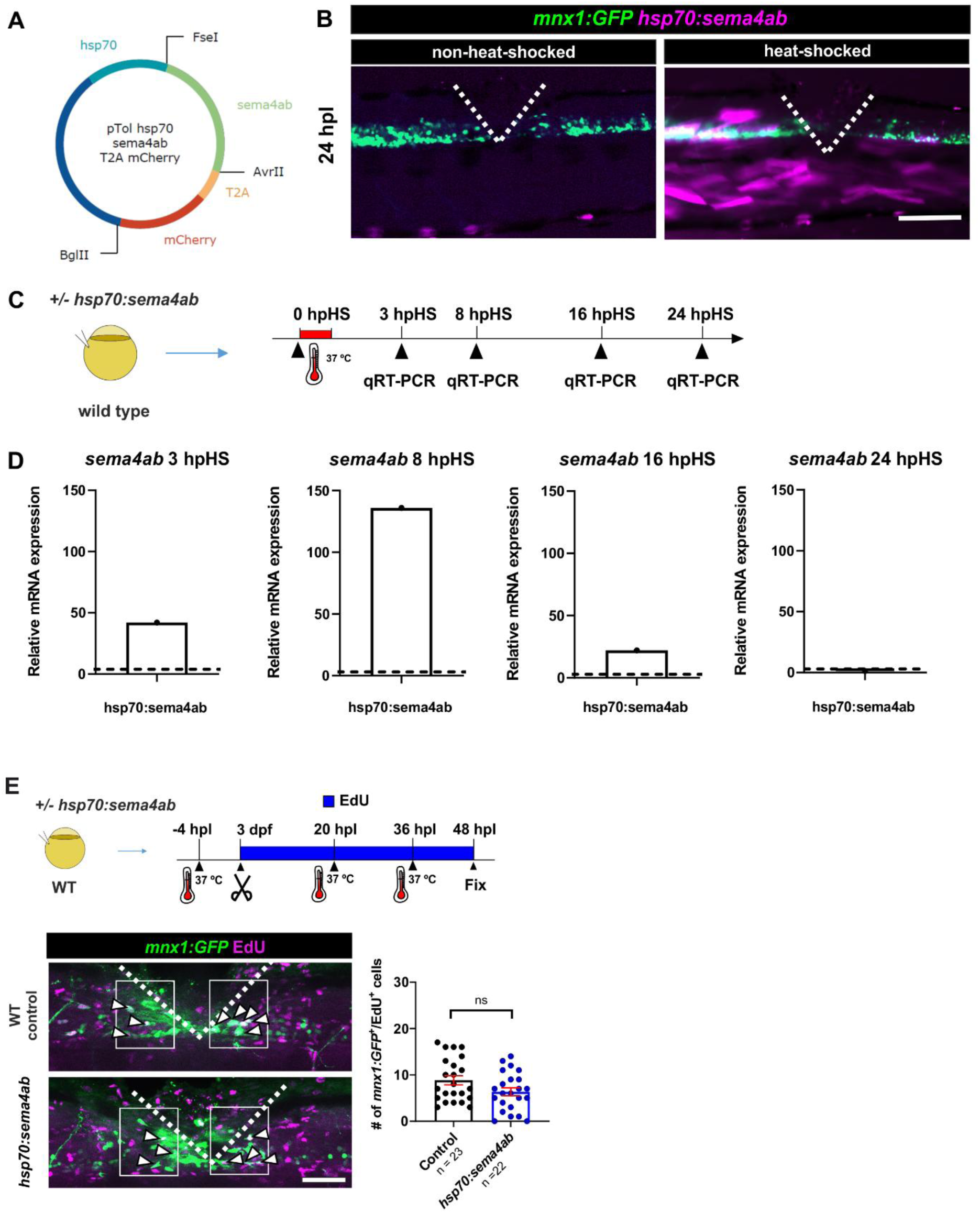
Efficient heat-shock overexpression of *sema4ab* does not alter regenerative neurogenesis in lesioned wild type larvae. **(A)** A map of the heat-shock vector injected to over-express *sema4ab* and a reporter (*mCherry*) is shown. **(B)** Heat-shock leads to detectability of mCherry protein mainly in muscle cells surrounding the injury site (position of ventral spinal cord indicated by *mnx1:GFP* transgene) at 24 hpl. **(C,D)** Experimental timeline to assess *sema4ab* over-expression by qRT-PCR after a single heat shock is shown in (C). qRT-PCR analysis shows that a single heat-shock leads to strongly increased *sema4ab* expression for at least 16 hours post heat-shock (hpHS, D). **(E)** Over-expression of *sema4ab* does not elicit any changes in the number of newly generated neurons after spinal lesion (F; gControl: 8.8 cells per larva ± 0.98; hsp70:sema4ab: 6.4 cells per larva ± 0.86; Mann-Whitney U-test: p = 0.3850). Error bars show SEM. Dotted lines show the injury site in B and E. Scale bars: 50 µm.

**Suppl. Fig. 6.**
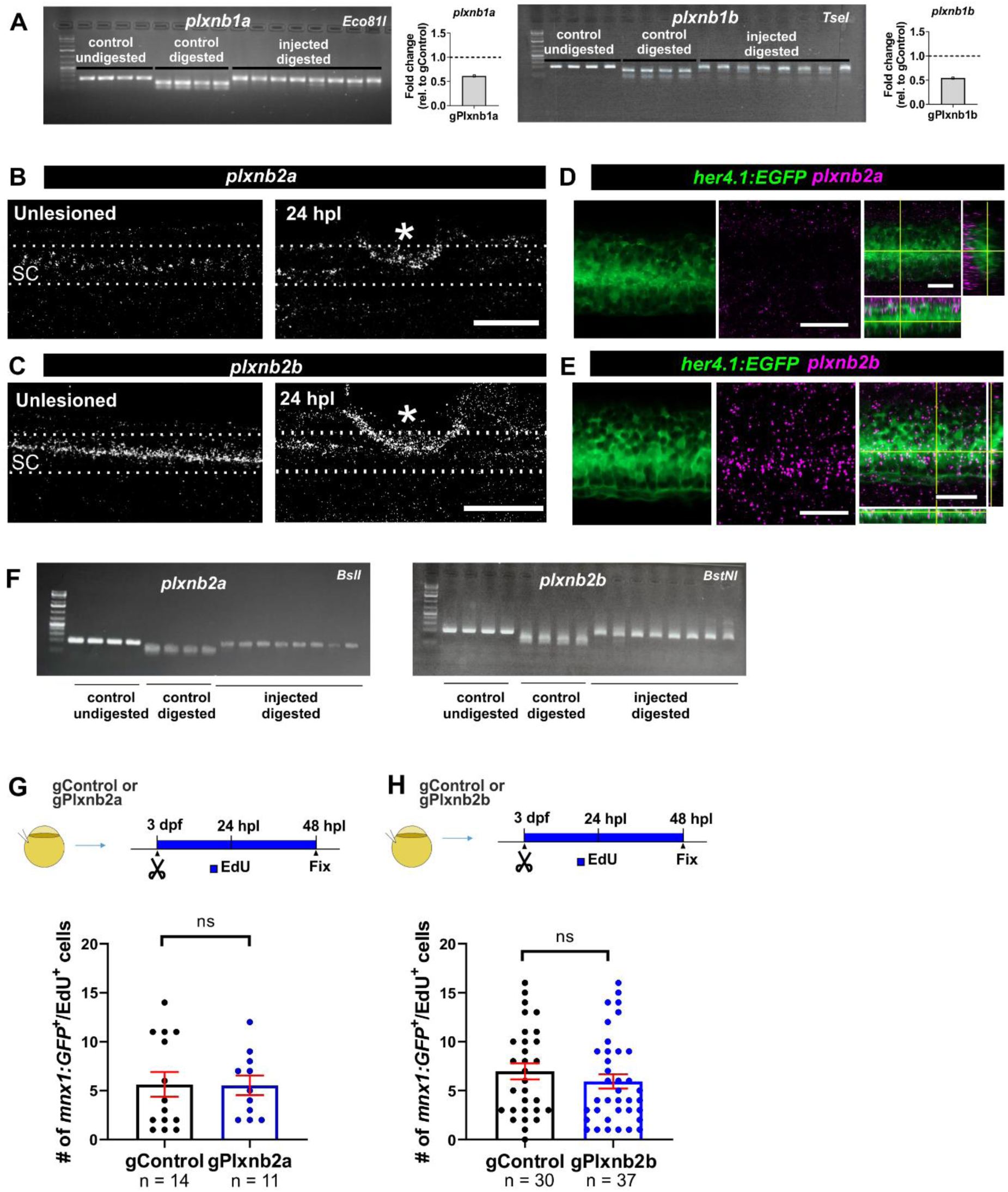
Somatic mutations of *plxnb2a and plxnb2b* do not affect regenerative neurogenesis. **(A)** RFLP and qPCR showing gRNA injection efficiency for *plxnb1a* and *plxnb1b*. For RFLP each lane represents one larva with and without digestion with the indicated restriction enzymes and with and without targeting these sites with gRNAs as indicated. Targeting the recognition sites with haCR gRNAs leads to efficient somatic mutation, as indicated by the almost complete resistance to digestion. Note a **∼**50% reduction in the RNA detected by qRT-PCR for either receptor. **(B,C)** HCR-FISH shows expression of *plxnb2a* (B) and *plxnb2b* (C) in a narrow domain in the spinal cord that does not change after lesion. Additional signal can be observed in the injury site after lesion for both genes (asterisks). **(D,E)** High magnifications of HCR-FISH for *plxnb2a* and *plxnb2b* show no detectable expression of *plxnb2a* at the level of ERGs (D), but of *plxnb2b* in *her4.1:EGFP*+ cells (E). **(F)** RFLP showing high gRNA injection efficiency for *plxnb2a* and *plxnb2b*. Each lane represents one larva with and without digestion with the indicated restriction enzymes and with and without targeting these sites with gRNAs as indicated. Targeting the recognition sites with haCR gRNAs leads to efficient somatic mutation, as indicated by the almost complete resistance to digestion. **(G,H)** haCR gene disruptions of *plxnb2a* (G, gControl: 5.6 cells per larva ± 1.26 ; gPlxnb2a: 5.5 cells per larva ± 1.00; Mann-Whitney U-test: p = 0.6938) or *plxnb2b* (H, gControl: 6.9 cells per larva ± 0.83 ; gPlxnb2b: 5.9 cells per larva ± 0.73; Mann-Whitney U-test: p = 0.3336) do not lead to changes in the number of newly generated motor neurons, compared to lesioned gControl-injected larvae. Error bars show SEM. Dotted lines delineate the position of the spinal cord. Scale bars: 100 µm (B, C), 25 µm (D, E).

**Suppl. Fig. 7.**
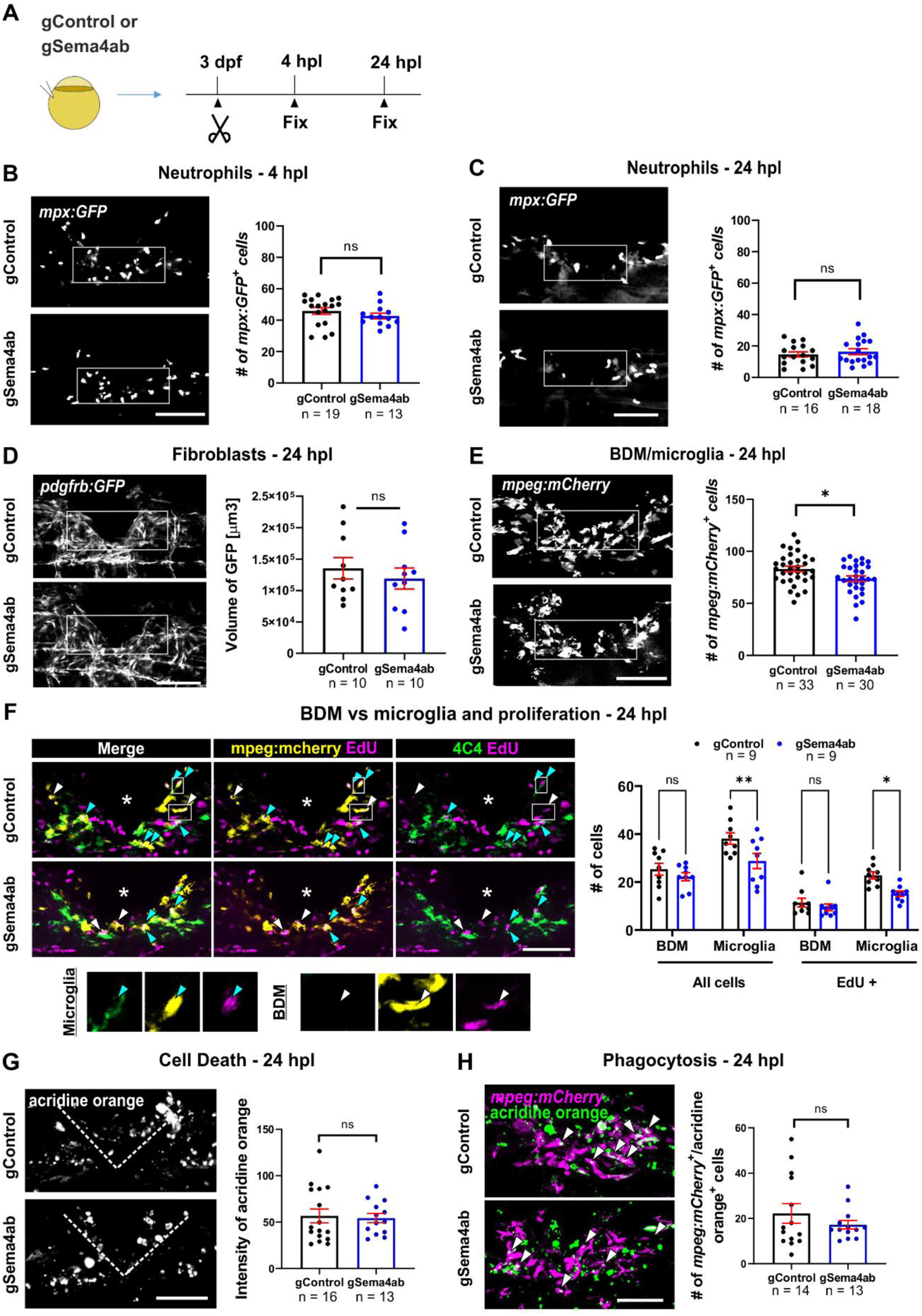
Cell recruitment and phagocytosis in the injury site are mostly comparable after *sema4ab* ablation. **(A)** A schematic indicating the experimental design for B-E is shown. **(B,C)** Neutrophil numbers do not show differences at 4 hpl (B, gControl: 45.89 cells per larva ± 2.1; gSema4ab: 42.69 cells per larva ± 1.8; Mann-Whitney U-test: p = 0.1854) and 24 hpl (C, gControl: 14.63 cells per larva ± 1.7; gSema4ab: 16.33 cells per larva ± 2.5; Mann-Whitney U-test: p = 0.5005) in *sema4ab* somatic mutants compared to control gRNA-injected larvae. **(D)** The abundance of fibroblast-like cells, measured by GFP-positive volume, shows no differences between controls and somatic mutants for *sema4ab* at 24 hpl (gControl: 135342 µm^3^ ± 17127; gSema4ab: 118911 µm^3^ ± 16774; Mann-Whitney U-test: p = 0.6842). **(E)** There is a reduction in the number of BDMs/microglia in gSema4ab compared to gControl larvae larvae at 24 hpl (gControl: 83.03 cells per larva ± 9.3; gSema4ab: 73.73 cells per larva ± 3.7; Mann-Whitney U-test: p= 0.0156). **(F)** There is a reduction in microglia cell number (gControl: 38.11 ± 2.3, Microglia 28.78 ± 3.2) but not for BDMs (gControl: 25.33 cells ± 2.5, gSema4ab: 22.22 cells ± 1.6) in *sema4ab* somatic mutants. The number of EdU-incorporating cells (incubation from lesion at 3 dpf to analysis at 24 hpl) is also reduced for microglia (gControl 22.8 ±1.4; gSema4ab 15.11 ±1.1, but not BDMs (gControl 11.4 ± 1.8; gSema4ab 9.4 ± 1.4; Šídák’s multiple comparisons test: BDM p =0 .7388; Microglia p = 0.0077, BDM/EdU p = 0.9328, Microglia/EdU p = 0.0392). **(G)** Cell death, measured by average labelling intensity of acridine orange in the lesion site, shows no differences between experimental groups (gControl: 56.67 ± 7.5; gSema4ab 54.27 ± 5.0; Mann-Whitney U-test: p = 0.8123). **(H)** BDMs/microglia phagocytosis, evaluated by counting *mpeg:mCherry+* cells with engulfed acridine orange+ particles, shows no differences between groups (gControl: 22.21 cells per larva ± 4.327; gSema4ab 17.23 cells per larva ± 1.912; Mann-Whitney U-test: p = 0.9714). White boxes designate quantification windows. Dotted lines outline the edges of the injury site. Error bars show SEM. Scale bars: 50 µm.

**Suppl. Fig. 8.**
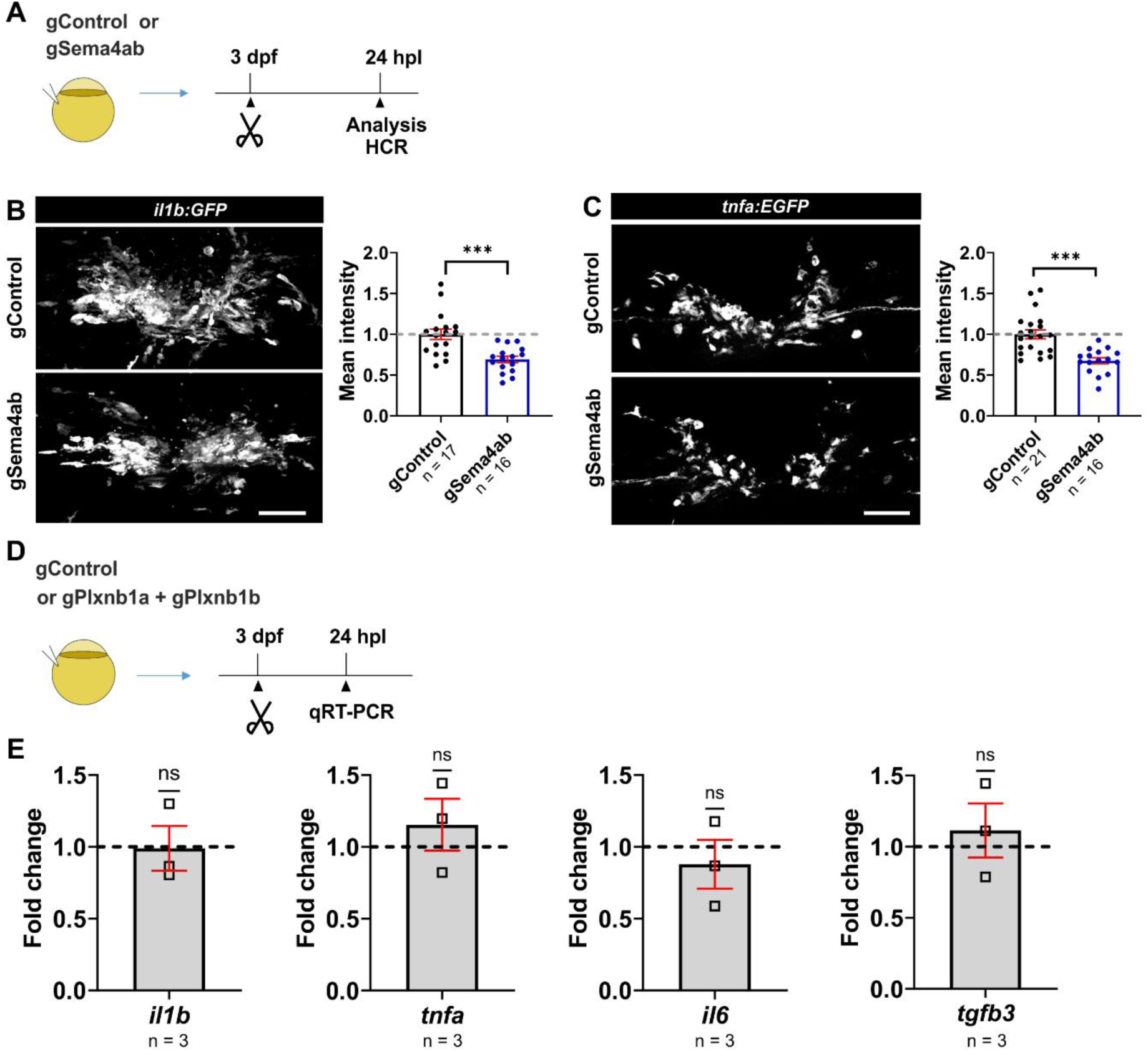
Loss of *sema4ab* reduces *il1b* and *tnfa* expression *in vivo, but plxnb1a/plxnb1b* disruption does not alter the cytokine profile of the lesion site. **(A)** A schematic indicating the experimental design for cytokine quantification *in situ* is shown. **(B)** *il1b:GFP* fluorescence intensity reported in transgenic fish is decreased in lesioned *sema4ab* somatic mutants compared to control gRNA-injected fish at 24 hpl (−1.45 fold-change; t-test: p = 0.0004). **(C)** *tnfa:EGFP* fluorescence intensity reported in transgenic animals is decreased in lesioned *sema4ab* somatic mutants compared to control injected fish at 24 hpl (−1.49 fold-change; t-test: p < 0.0001).**(D)** A schematic indicating the experimental design for qRT-PCR. **(E)** qRT-PCR analyses of major pro-inflammatory cytokines (*il1b, tnfa, il6*) and *tgfb3* in *plxnb1a/plxnb1b* double somatic mutants show no differences to injured control gRNA-injected larvae at 24 hpl (One-sample t-test; *il1b*: p = 0.537; *tnfa*: p = 0.4840; *il6*: p = 0.5464; *tgfb3*: p = 0.687). Error bars show SEM. Scale bars: 50 µm.

**Suppl. Fig. 9.**
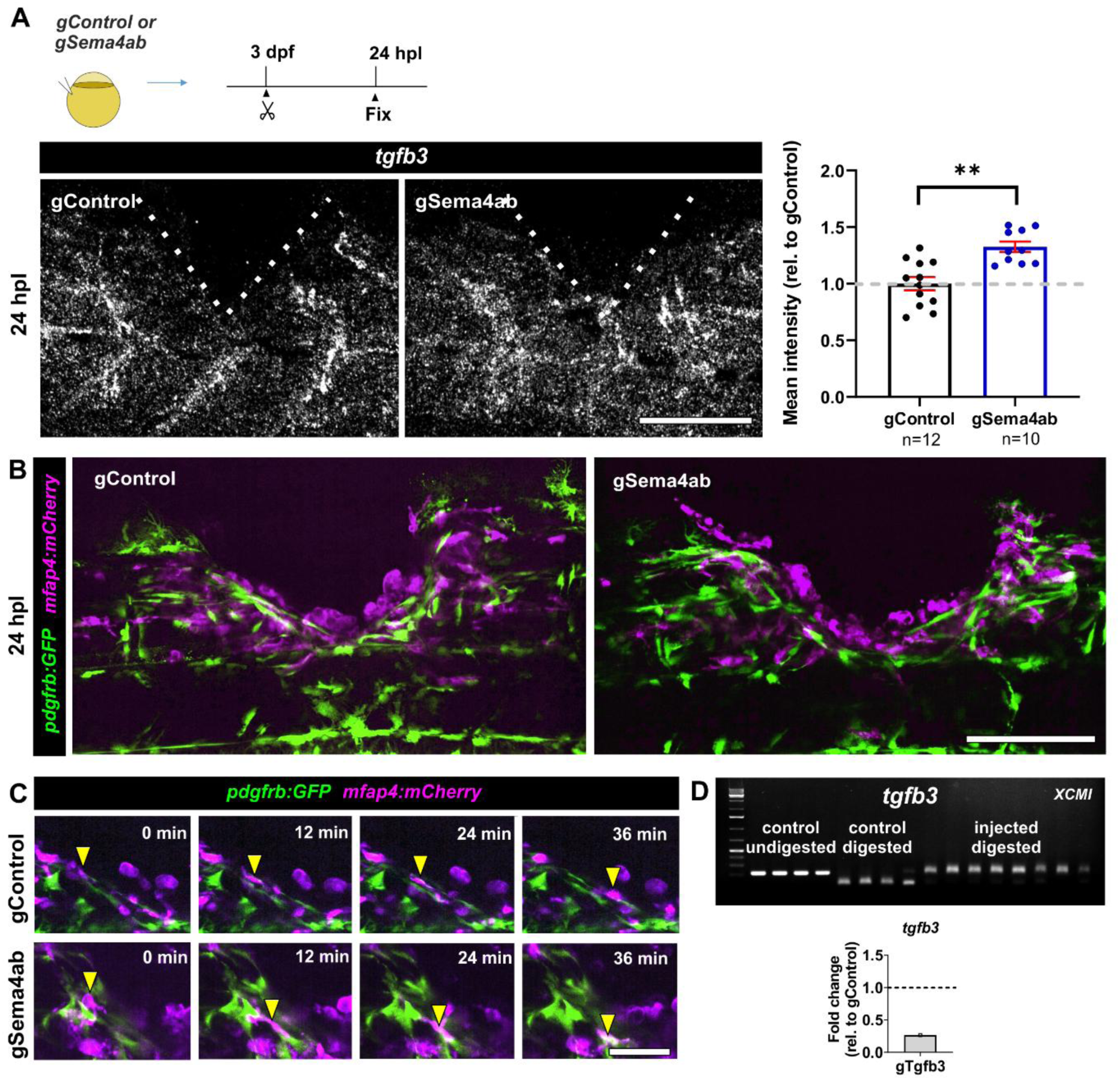
Experimental manipulations and cell interactions in the lesion. **(A)** HCR-FISH for *tgfb3* shows an increase in fluorescence intensity in lesioned *sema4ab* somatic mutants compared to lesioned control gRNA-injected fish at 24 hpl (gControl: 1.0 ± 0.058; gSema4ab: 1.3 ± 0.046; t-test: p = 0.0011). Dotted lines show the injury site. **(B)** Individual frames of time-lapse movies show *mfap4:mCherry*+ cells in close contact to *pdgfrb:GFP*+ cells in both control gRNA-injected animals and in *sema4ab* somatic mutants. **(C)** Time series of movie frames show that *mfap4:mCherry+* cells (arrowheads) migrate along fibroblast processes in both conditions**. (D)** RFLP and qPCR showing gRNA injection efficiency for *tgfb3*. For RFLP each lane represents one larva with and without digestion with the indicated restriction enzymes and with and without targeting these sites with gRNAs as indicated. Targeting the recognition sites with haCR gRNAs leads to efficient somatic mutation, as indicated by the almost complete resistance to digestion. Note a **∼**50% decay in the RNA detected by qPCR. Scale bars: 100 µm (A, B,), 15 µm (C).

**Suppl. Fig. 10.**
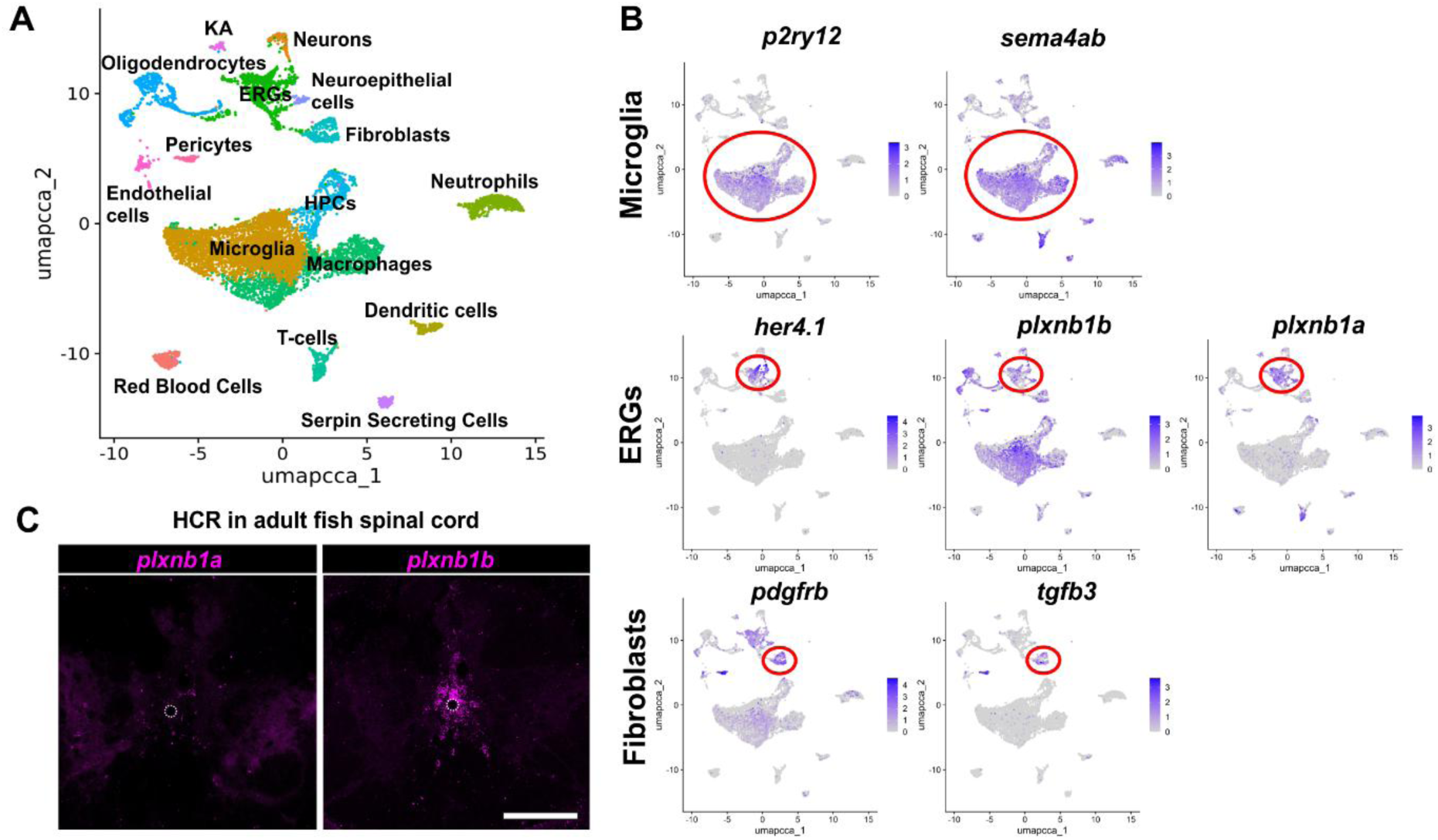
scRNA-seq shows that genes analysed in this study are also expressed in a spinal injury site in adult zebrafish. **(A)** UMAP of the main cell types in the lesion site of adult zebrafish from Cigliola et al. 2023 *Nat Commun* 14:4857 is shown. **(B)** Feature plots show the expression of reference genes for relevant cell types (microglia: *p2ry12*; ERGs: *her4.1*; and fibroblasts-like cells: *pdgfrb*), as well as the expression of genes of interest for this study, *sema4ab, plxnb1a, plxnb1b* and *tgfb3*. Circles label expression in cell types that is conserved between larvae and adults. **(C)** HCR-FISH in cross sections of unlesioned spinal cords of 4-month-old adult zebrafish show expression of *plxnb1a and plxnb1b* in the ventricular zone. Dorsal is up and ventral is down. Dotted circle shows the central canal. Scale bar: 50 µm.

**Suppl. Fig. 11.**
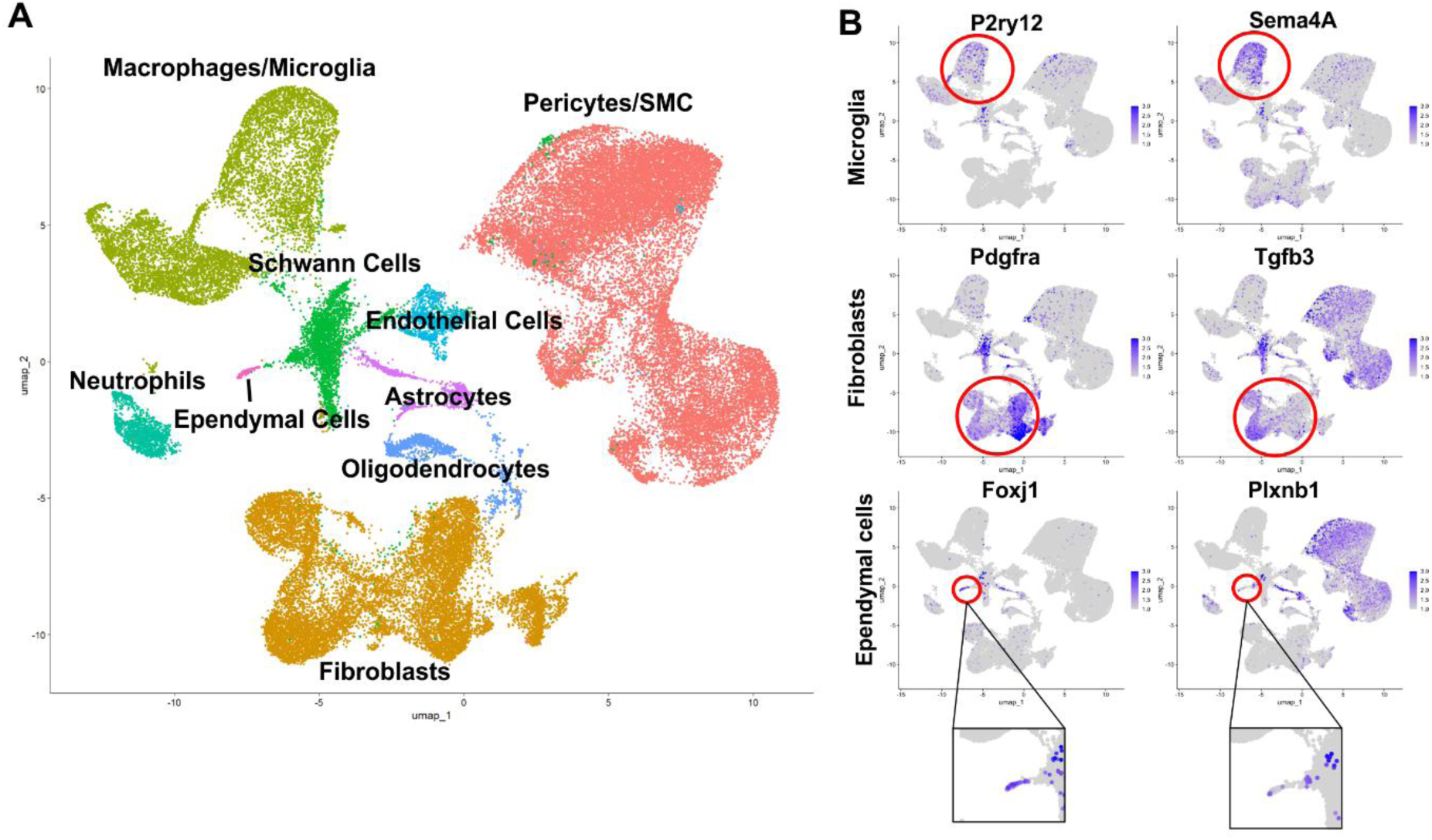
scRNA-seq shows that genes analysed in this study are also expressed in a spinal injury site in adult mice. **(A)** A UMAP representation indicating the main cell types in a mouse spinal lesion from Xue et al., 2024 *Nat Commun* 15:6321 is shown. **(B)** Expression pattern of mouse ortholog genes *Sema4A*, *Tgfb3* and *Plxnb1* is conserved in microglia (*P2ry12*), fibroblasts (*Pdgfra*) and ependymal cells (*Foxj1*), respectively. Circles label expression in cell types that is conserved between zebrafish and mouse.

## SUPPLEMENTARY TABLES

**Suppl. Table 1:**
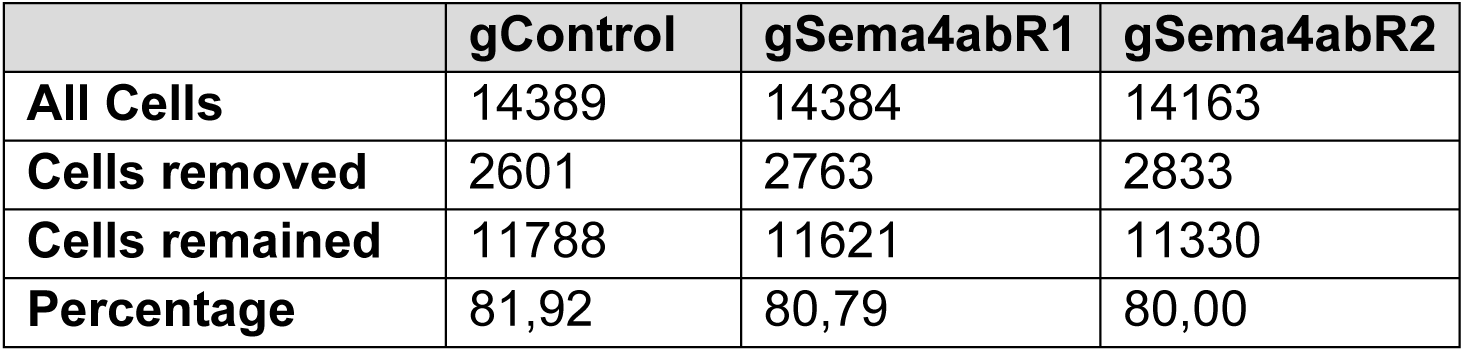
Cell yields of scRNA-seq experiments.

|  | **gControl** | **gSema4abR1** | **gSema4abR2** |
| --- | --- | --- | --- |
| **All Cells** | 14389 | 14384 | 14163 |
| **Cells removed** | 2601 | 2763 | 2833 |
| **Cells remained** | 11788 | 11621 | 11330 |
| **Percentage** | 81,92 | 80,79 | 80,00 |

**Suppl. Table 2:**
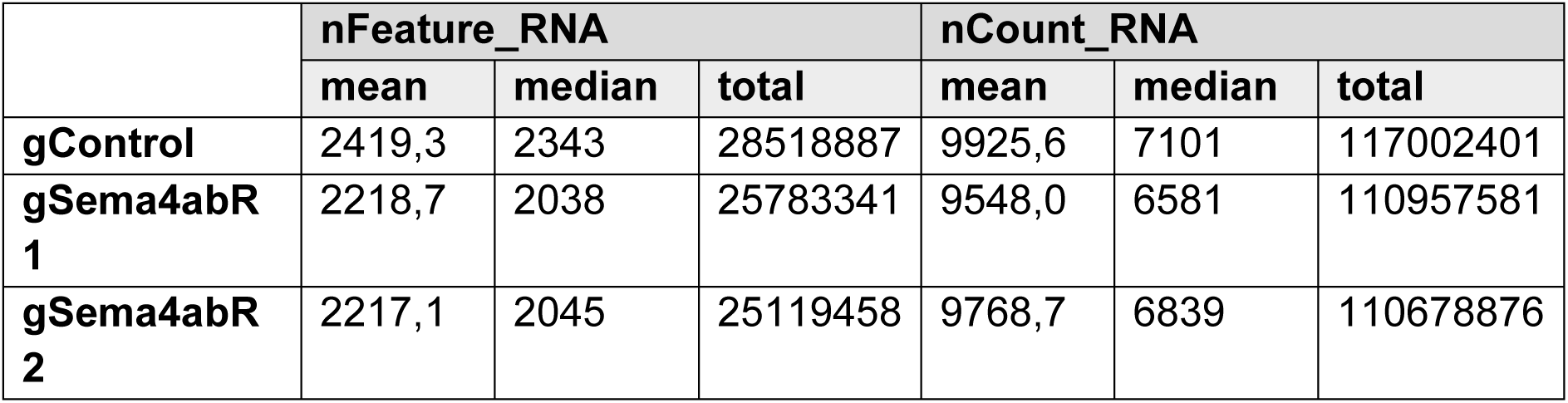
The mean, median and total number of reads (nCount_RNA) and unique genes (nFeatures_RNA) detected per sample.

**Suppl. Table 3:** The number of reads mapped to the *sema4ab* gene and the number of reads mapped to the *sema4ab* haCR-targeted site.

|  | ***sema4ab* mapped total reads** | **target site total reads** | **target site intact reads** |
| --- | --- | --- | --- |
| **gControl** | 16945 | 151 | 151 |
| **gSema4abR 1** | 9880 | 4 | 0 |
| **gSema4abR 2** | 10384 | 6 | 0 |

**Suppl. Table 4:**
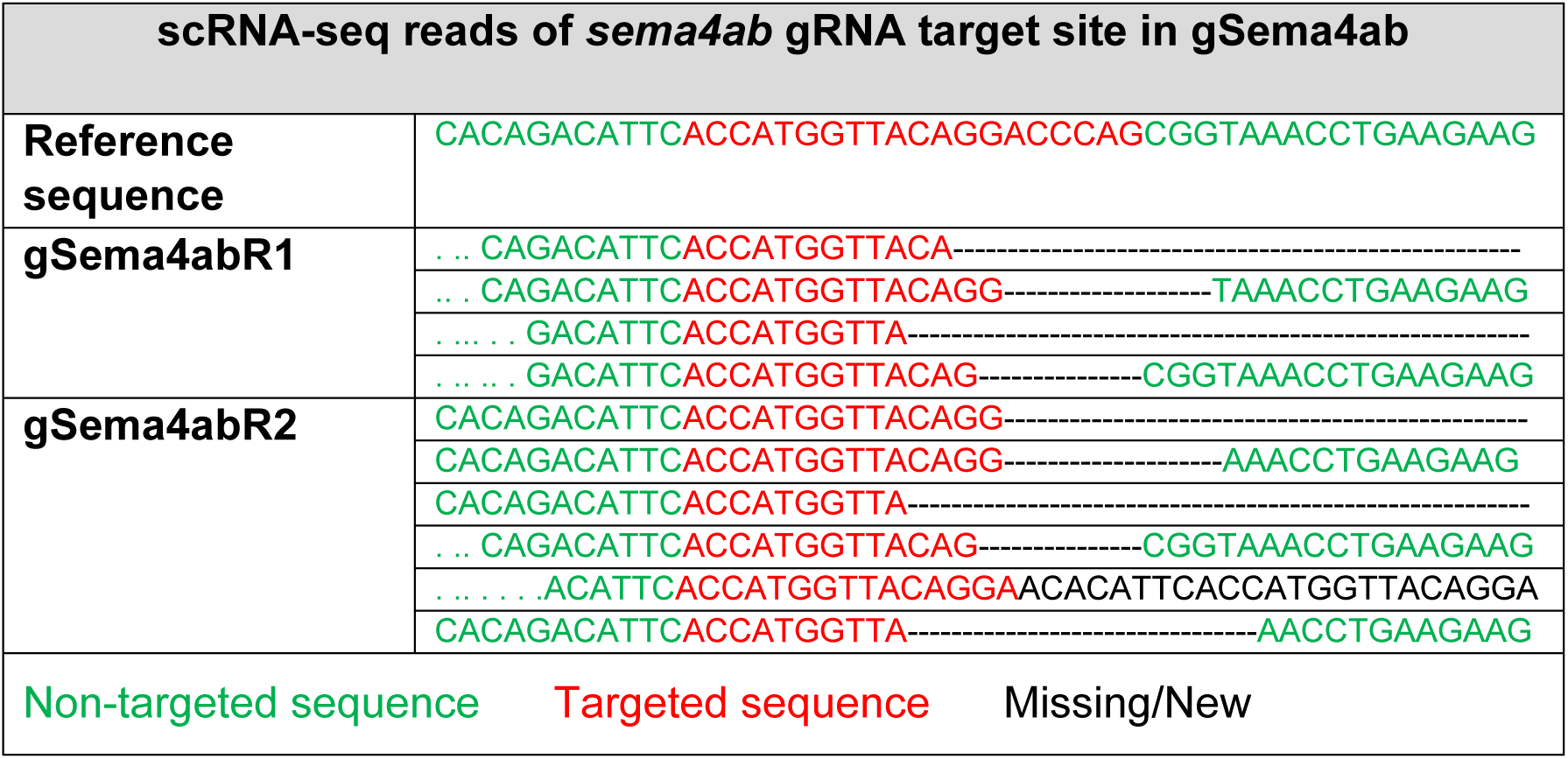
Examples of reads that span the haCR target site in the *sema4ab* haCR-injected samples in scRNA-seq experiments.

**Suppl. Table 5.**
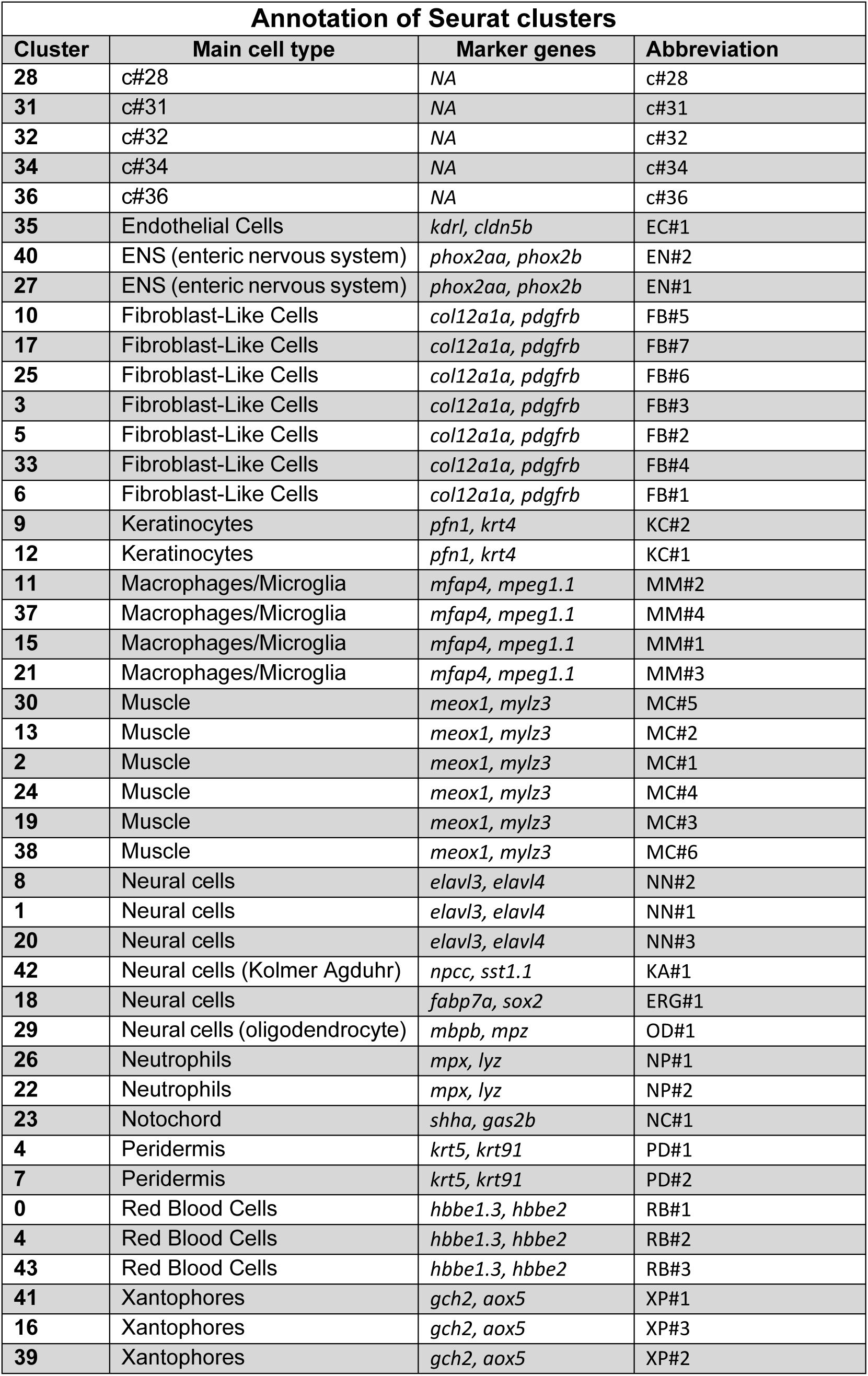
Cell cluster annotation reference. The table shows the correspondence between each cluster obtained by unsupervised clustering and the correspondent main cell type and cluster name assigned.

**Suppl. Table 6:**
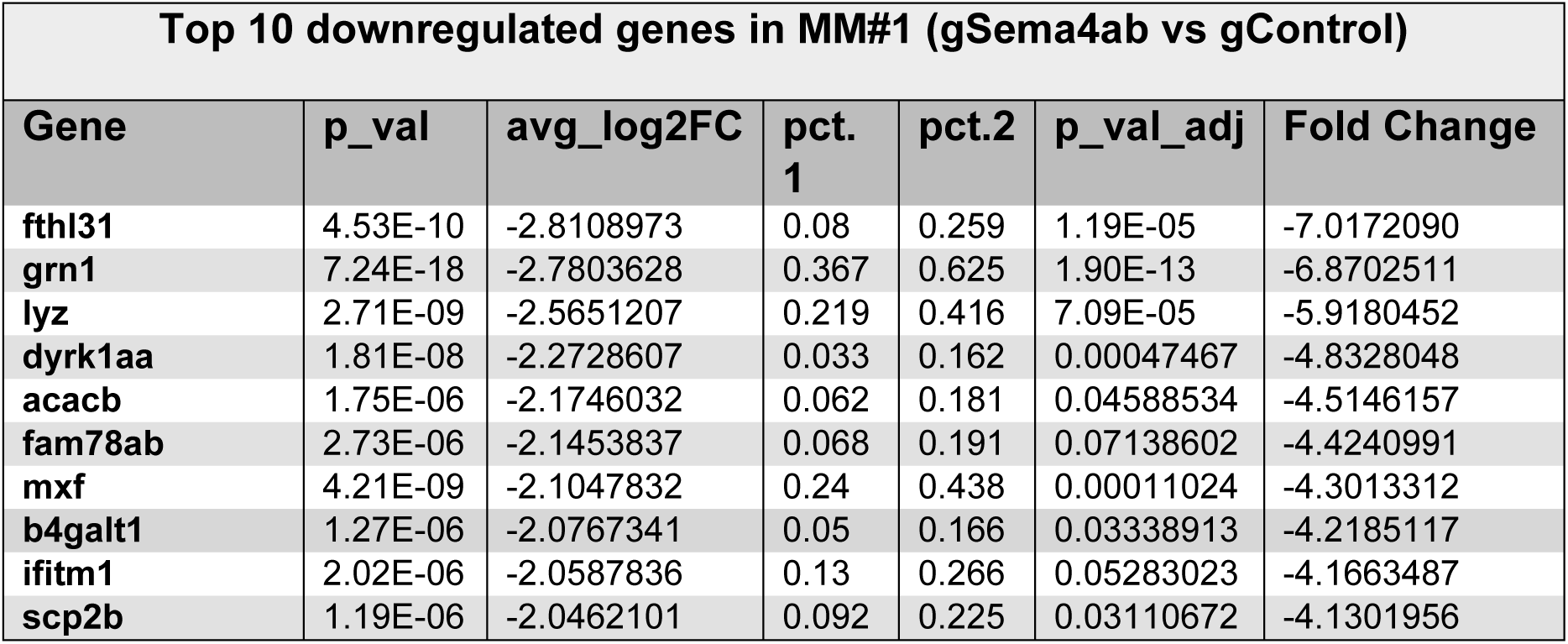
Top 10 downregulated genes in MM#1 after *sema4ab* somatic mutation in scRNA-seq.

**Suppl. Table 7:**
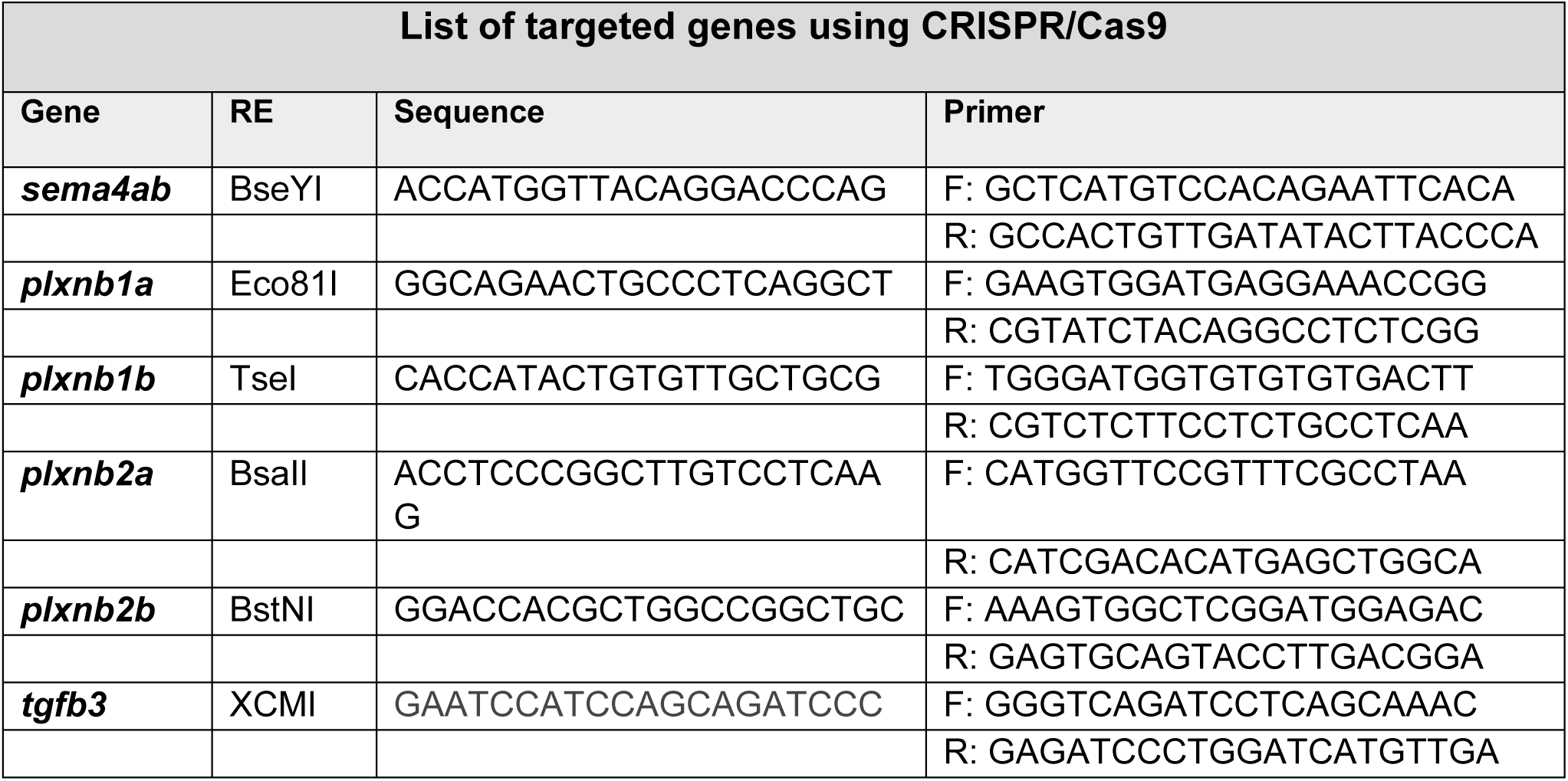
Table showing the gene sequences targeted with gRNAs. Each column represents the targeted sequence of a gene and the primers and restriction enzymes used to validate the efficiency of the mutation.

**Suppl. Table 8:** Primers used for cloning the *mfap4:mCherry-CAAX;myl7:mCerulean* vector.

| **Primers use for cloning the *mfap4:mCherry-CAAX;myl7:mCerulean* vector** |  |
| --- | --- |
| **Primer** | **Primer sequence (5'-3')** |
| *mfap4* | **Fw:** ttt ttt gag ctc ctc gag gcg ttt ctt ggt ac |
|  | **Rev:** ttt ttt ggc gcg cct gga tcc cac gat cta aag tca tga ag |
| *mCherry-CAAX* | **Fw:** aaa aaa gaa ttc gcc acc atg gtg agc aag ggc gag g |
|  | **Rev:** aaa aaa tta att aat cag gag agc aca cac ttg cag ctc atg cag ccg ggg cca ctc tca tca gga ggg ttc agc tta gat ctg agt ccg gac ttg tac agc tcg tcc atg ccg aga gtg |

**Suppl. Table 9.**
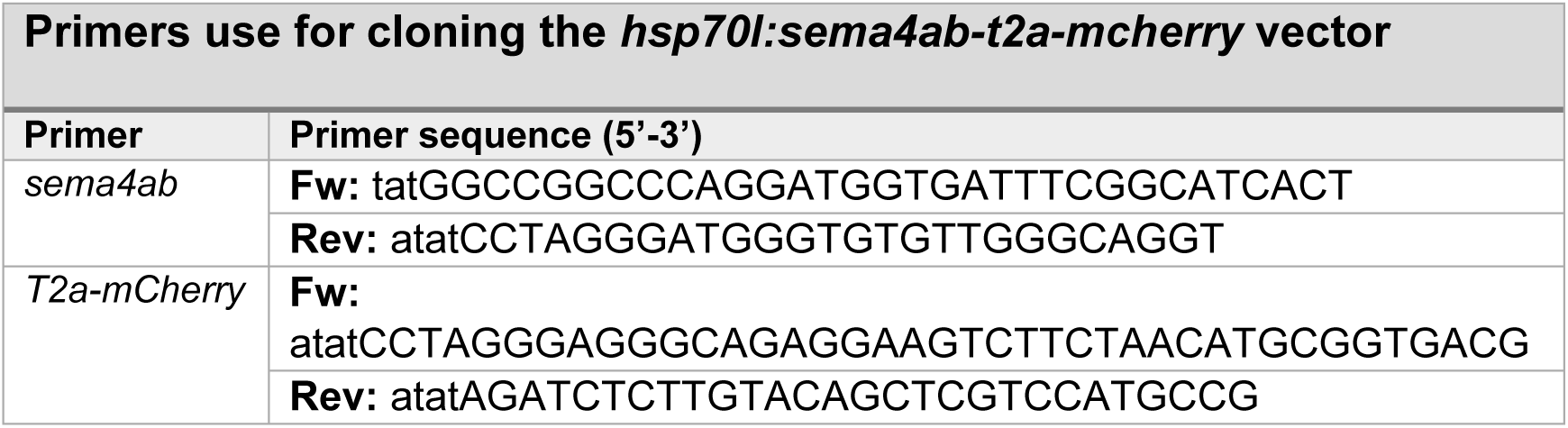
Primers used for cloning the *hsp70l:sema4ab-t2a-mCherry* vector.

**Suppl. Table 10:**
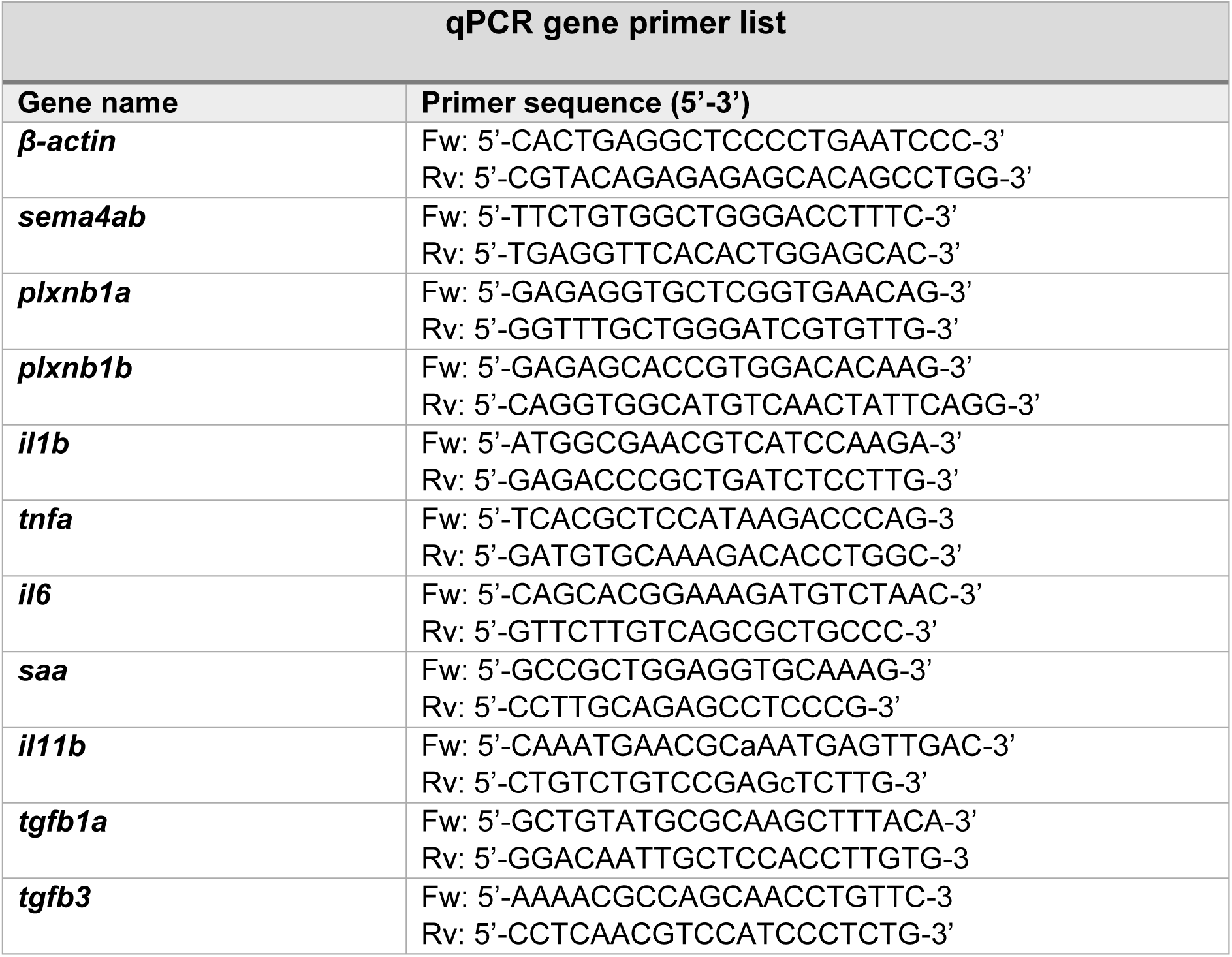
List of qRT-PCR primers.

## Supplemental data files

<u>Supplemental data file 1:</u> Changes in transcript abundance for macrophage activation pathways detected by KEGG analysis. Note that most genes are down-regulated.

<u>Supplemental movie 1</u>: lesioned control-gRNA injected

<u>Supplemental movie 2:</u> lesioned *sema4ab* haCR-injected. Both movies run from 10 to 25 hpl (6 min intervals).

